# Discovery of alternative stable states in a synthetic human gut microbial community

**DOI:** 10.1101/2024.11.28.625814

**Authors:** Daniel Garza, Bin Liu, Charlotte van de Velde, Pallabita Saha, Xingjian Zhou, Didier Gonze, Kenneth Simoens, Kristel Bernaerts, Karoline Faust

**Affiliations:** Department of Microbiology, Immunology and Transplantation, Rega Institute for Medical Research, Laboratory of Molecular Bacteriology, KU Leuven, Leuven B-3000, Belgium; Université Paris-Saclay, INRAE, PROSE, 92160 Antony, France; Key Laboratory of Environmental Biotechnology, Research Center for Eco-Environmental Sciences, Chinese Academy of Sciences, Beijing, People’s Republic of China; Unité de Chronobiologie Théorique, Faculté des Sciences, CP 231, Université Libre de Bruxelles, Bvd du Triomphe, B-1050 Bruxelles, Belgium; Department of Chemical Engineering, Chemical and Biochemical Reactor Engineering and Safety (CREaS), KU Leuven, B-3001 Leuven, Belgium

## Abstract

Several human-associated microbial communities exist in multiple configurations and can change their composition in response to perturbations, remaining in an altered state even after the perturbation ends. Multistability has been previously proposed to explain this behavior for gut microbiota in particular, but has not been clearly demonstrated experimentally. Here, we first investigated the life history strategies of three common human gut bacteria to identify mechanisms driving alternative states. We then used this data to build and parameterize a kinetic model, which predicted that alternative states emerge due to phenotype switching between subpopulations of the same species. Perturbation experiments supported these predictions, and confirmed the existence of alternative states. Finally, simulations showed that phenotype switching can also explain alternative states in larger communities. Thus, a transient perturbation combined with metabolic flexibility is sufficient for alternative communities to emerge, implying that they are not necessarily explained by differences between individuals.

**One-Sentence Summary:** We demonstrate the existence of alternative states in a human gut microbial community and propose phenotype switching as a mechanism explaining their emergence.

## Main Text

Several human-associated microbial communities assemble into more than one configuration^1–3^ and change their composition in response to perturbations, remaining in an altered state even after the perturbation ends^4,5^. While various hypotheses have been proposed to explain this behavior^6–8^, clear demonstrations of the mechanisms underlying these hypotheses are still lacking. One common explanation, supported by empirical evidence, suggests that microbiomes, like many ecological systems, can assemble into alternative stable states^7^. Thus, even when the same microbes are assembled under similar environmental conditions, they may converge on distinct community compositions, influenced by their assembly history^9^. Additionally, minor yet continuous changes in environmental parameters could lead to significant shifts in community states^10^. Mechanistically identifying alternative stable states and regime shifts in natural communities has historically been challenging^11^. This is because organisms can change their own environments—microbes, for example, consume resources^12^, change the environment’s pH^13^, and create physical structures^14^—in response to biotic and abiotic changes, thereby making it difficult to disentangle community states from environmental factors.

Microbial cells exist in dynamic equilibrium, coexisting with other cells and their environments^15^. Their metabolic capabilities are encoded in their genomes, but the metabolic programs they execute depend on the differential expression of enzymes^16^. This differential expression enables a variety of metabolic strategies, which are evolvable and can be flexible, heterogeneous, and dynamic^17–19^. For instance, when exposed to a mix of substrates, cells might use these substrates either simultaneously or sequentially^20^—first consuming one and then another—or they might alternate between both strategies^21^.

While the dynamic metabolic strategies of microbes, gene regulation, and phenotype switching have been extensively studied in isolates since Jacques Monod’s seminal work^22^, their impact on microbiome ecology and stability remains vastly underexplored^23^. In our previous study, we observed that the ecological interactions between two gut bacterial species changed during co-culture^24^. These changes were in response to alterations in pH and the concentration of degradation products resulting from their metabolic strategies. Given that bacteria express alternative metabolic programs under varying environmental conditions, we hypothesize that the coexistence and switching between bacterial growth strategies could induce sharp transitions in community-level phenotypes, leading to multistability and the emergence of predictable alternative community states.

## Life history strategies of gut bacteria in a heterogeneous environment

To test this hypothesis, we used a simple three-species gut community that allowed us to combine in vitro experiments with mechanistic modeling. We first investigated the individual phenotypes of three common human gut bacteria in Wilkins-Chalgren (WC) anaerobic medium. This medium includes two simple carbon sources, glucose and pyruvate, along with substrates from tryptone and yeast extract, which notably contain measurable amounts of trehalose (average 0.71 mM +/-0.07). The composition of this medium is simple enough to allow us to track the kinetics of key metabolites, yet its complex components mimic the nutrient heterogeneity expected in the colon.

We used genome-scale metabolic models to derive sets of biochemical reactions that define the core energy metabolism of each species. We then collected RNA-seq data at different growth stages to confirm the activities of these pathways (Fig. 1 A-C). These core pathways connect the import of carbon sources with the production of fermentation acids, enabling us to compare model predictions with the measured data (Fig. 1 D-F). By analyzing these pathways alongside live cell growth kinetics, medium pH changes, and metabolite composition, we were able to outline bacterial life history strategies^25^. We incorporated these strategies into an ordinary differential equation model (Supplementary Text S1). This model was calibrated against experimental data, as indicated by the traced lines in Fig. 1 D-F.

**Fig. 1:**
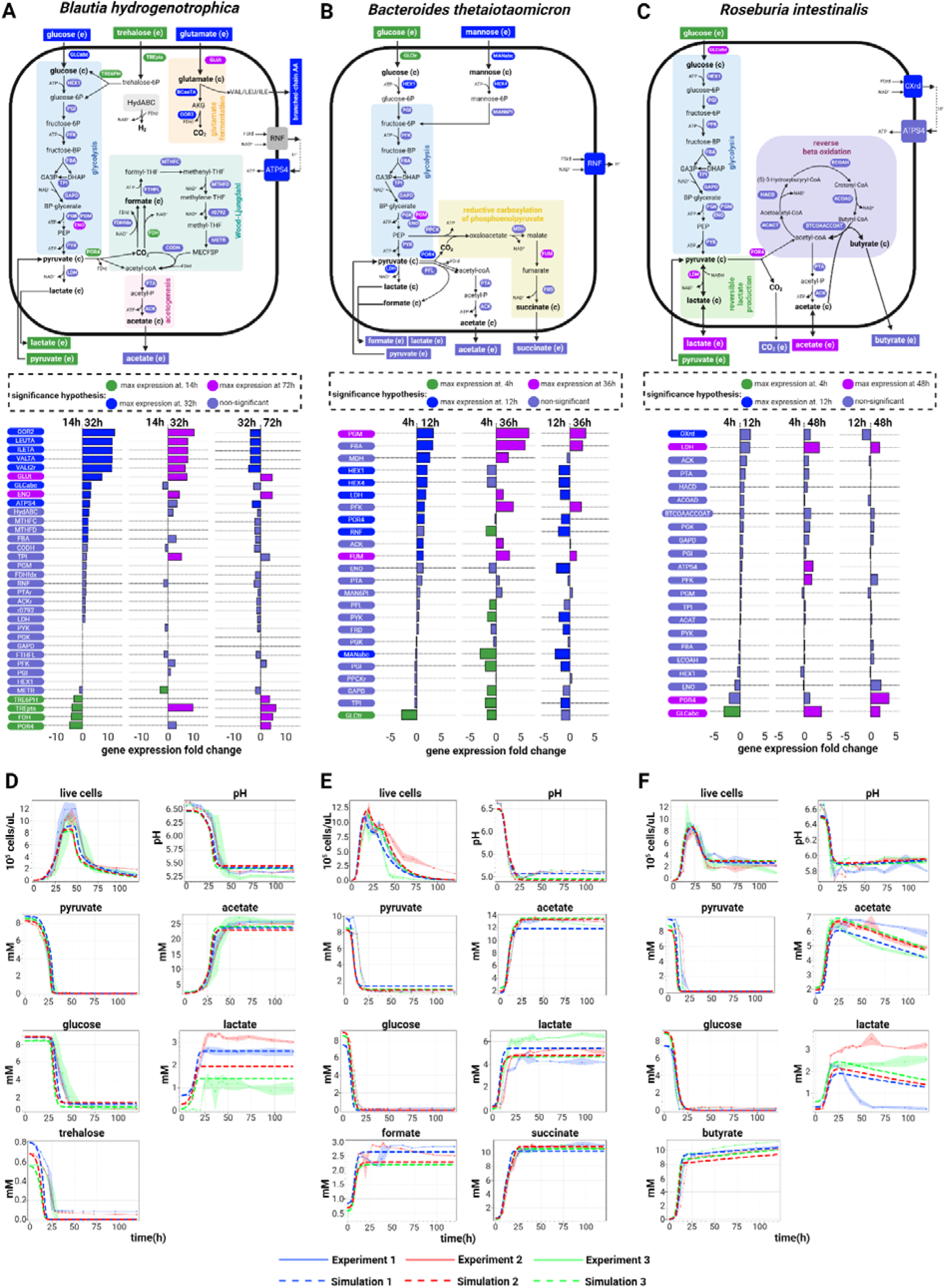
Growth kinetics and modeled metabolism of three human gut bacteria. Schematic illustration of the core energy metabolism pathways for *Blautia hydrogenotrophica* (**A**), *Bacteroides thetaiotaomicron* (**B**), and *Roseburia intestinalis* (**C**), as deduced from genomic metabolic model reconstructions coupled with RNA-seq data for temporal pathway activity, and empirical growth data. The middle panel details gene expression changes observed during cultivation, with corresponding gene names linked to their associated reactions in the upper panel. The hypothesis tests were performed using DESeq2. The Figure also depicts the consumption of carbon sources (glucose, pyruvate, and trehalose) and the generation of fermentation products (including acetate, lactate, and butyrate). Panels (**D**-**F**) compare experimental growth data over time with model simulations (represented by dashed lines) for the three species cultured in WC medium. The growth data represent averages from three independent monoculture experiments, each with 3-6 biological replicates (solid dots are the averages and the shaded lines reflect the standard deviations of biological replicates). The simulation’s initial conditions were the same as the experimental setups.

Briefly, under our growth conditions, *Blautia hydrogenotrophica* initially consumes trehalose via a trehalose-specific PTS transporter. The gene for this transporter (*TREpts*) is overexpressed in the early stages of growth compared to the later stages (Fig. 1A). It switches to glucose utilization only after trehalose is depleted, facilitated by a non-PTS glucose transporter (*GLCabc*) that is inhibited during trehalose consumption (Fig. 1A). Interestingly, its genome lacks the glucose-specific IIA component gene of the PTS system, commonly found in closely-related *Blautia* and *Ruminococcus* strains (Supplementary Table S1). We confirmed this sequential substrate preference by showing that higher trehalose concentrations extended *Blautia hydrogenotrophica*’s non-glucose consuming phase (Supplementary Fig. S1). Flow cytometry showed a clear bimodal population distribution during this transition, suggesting the presence of similar subpopulation sizes of non-dividing trehalose consumers and dividing glucose consumers (Supplementary Movie S1). The growth rate increased during glucose consumption compared to the trehalose phase (Fig. 1D, ∼26 hrs). Equations modeling *Blautia hydrogenotrophica*’s life history strategy are detailed in Supplementary Text S1.

*Bacteroides thetaiotaomicron* rapidly metabolizes glucose and pyruvate, producing fermentation acids that significantly reduce the medium’s pH (Fig. 1E). However, this organism is inhibited at low pH conditions (i.e., pH<5.5)^26^. When the carbon sources are exhausted, most cells lose viability at low pH but can still be detected through flow cytometry. The loss of membrane integrity was confirmed using propidium iodide staining. To reflect this in our model, we introduced functions that describe transitions from active to inactive subpopulations, triggered by nutrient scarcity in acidic environments (see Supplementary Text S1). We consistently observed a second growth peak before a major population inactivation (as shown in Fig. 1E), which we believe is due to trace mannose consumption, consistent with our previous findings^24^. Mannose depletion was verified through measurements and gene expression analysis, although the precise kinetics remain unresolved.

*Roseburia intestinalis* generates butyrate through the reverse β-oxidation pathway (illustrated in Fig. 1C). In our experiments, *R. intestinalis* efficiently consumed glucose and pyruvate, producing butyrate, acetate, and lactate, impacting the pH to a lesser extent than *B. thetaiotaomicron*. As we previously described^24^, in the absence of glucose, *R. intestinalis* transitions to a slow growth mode, characterized by the prolonged survival of viable cells (evident in “live cells” curves in Fig. 1F), sporadic consumption of lactate/acetate (Fig. 1F), and continuous butyrate production (also in Fig. 1F). In our model, we represented this average behavior by incorporating rapid cell death in the absence of glucose, and by shifting subpopulations to slow growth in response to lactate and acetate (detailed in Supplementary Text S1). However, our model does not fully account for the observed heterogeneous lactate utilization across different experiments (as depicted in Fig. 1F).

## The model’s stability landscape reveals sharp transition zones

After calibrating our model with monoculture growth data (Fig. 1 D-F and Supplementary Text S1) and validating its performance in batch cocultures (Supplementary Fig. S2), we incorporated dilution terms to simulate in silico the stability landscape of the community in a continuous culture environment. We explored how steady-state concentrations of bacteria and metabolites respond to controllable factors—medium pH and dilution rate—which do not directly alter their initial concentrations but can impact the system’s dynamics. The dilution rate impacts the steady-state concentration of metabolites and nutrient availability, thereby affecting population growth rates and the transition between metabolic phenotypes (Fig. 2A), while pH directly impacts growth. Changes in growth also impact the production of fermentation acids and, for instance, their subsequent utilization by *Roseburia intestinalis*’ slow growth mode. In summary, these parameters significantly shape the overall community phenotype (Fig. 2B and 2C).

**Fig. 2:**
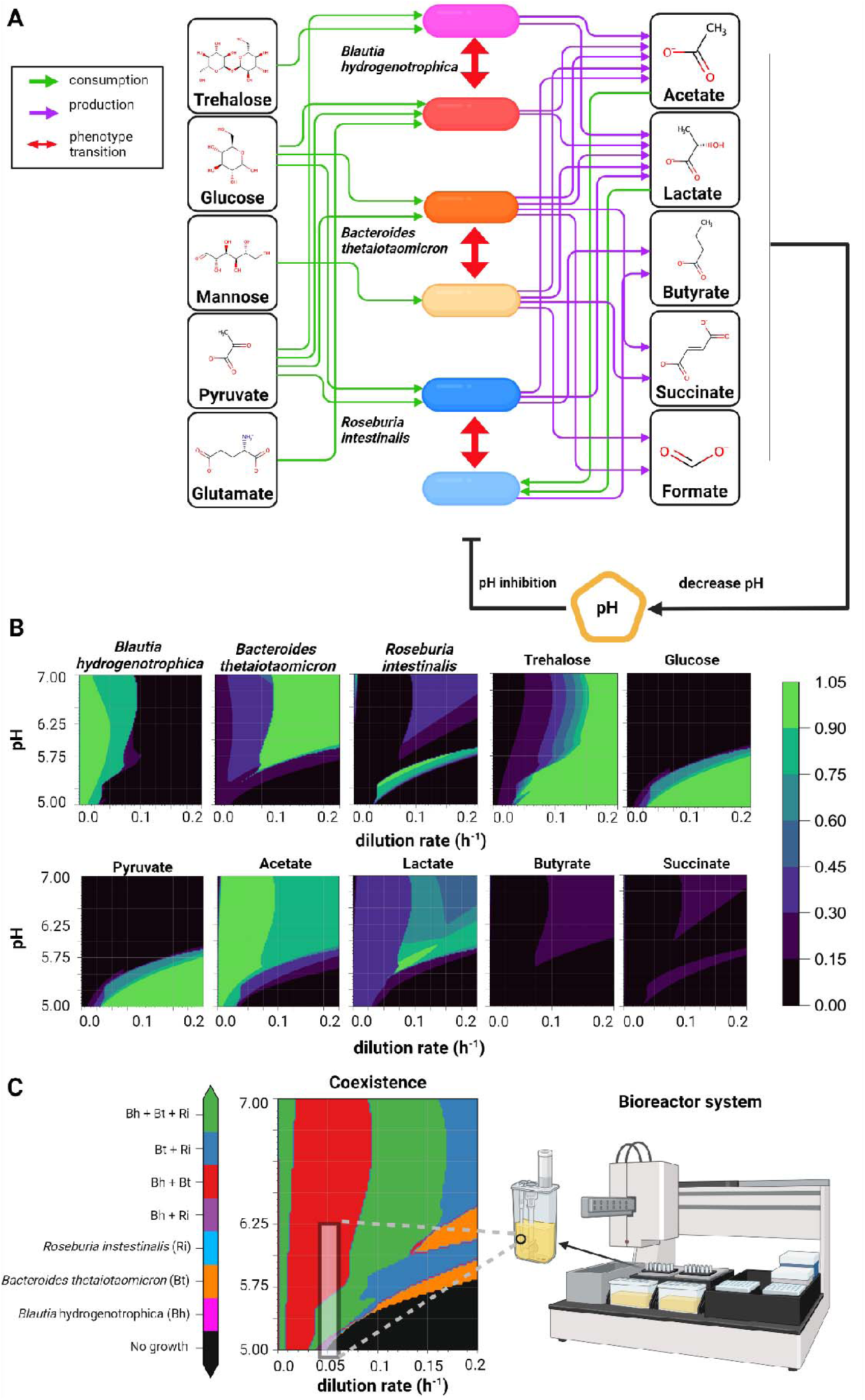
Mechanistic modeling reveals that pH and dilution rate drive community transitions towards alternative stable states. (**A**) Schematic of the mechanistic model encoded as ordinary differential equations (see Supplementary Text S1). The model incorporates experimental data (refer to Fig.1B-D and Supplementary Figs.S1 and S2) to simulate variable conditions. (**B**) Contour plots depicting the predicted steady-state concentrations of model variables across a pH range of 5.0-7.0 and dilution rates from 0.0 to 0.2 h^-1^. Concentrations, initially in millimolar (mM) for metabolites and in 10^5^ cells/µL for cells, are normalized to their maximum values for comparative purposes. (**C**) A community composition phase plot indicates the presence or absence of species across different pH and dilution rate conditions. The black shaded rectangle in the plot highlights the conditions that are within the range observed in our minibioreactor experiments (refer to Figs. 3 and 4).

A surprising observation that resulted from this in silico analysis of the community is that the stability landscape of some of the state variables revealed sharp transitions between steady-state metabolite and bacterial cell concentration profiles. For example, *Blautia hydrogenotrophica* maintains consistent concentrations under wide pH ranges, but undergoes a sharp shift when the dilution rate increases beyond a certain level (Fig. 2B). *Roseburia intestinalis* survives in two separate zones, including a narrow range of conditions where it outcompetes the other species (Fig. 2C). These zones are separated by areas where no growth occurs. Overall, the presence of sharp transitions between zones in the stability landscapes suggests that alternative stable states can emerge depending on the initial concentrations. It also indicates the presence of tipping points, where minor environmental changes can lead to significant shifts in community structure.

## Phenotype-switching explains multistability in silico and agrees with observations in vitro

To understand the mechanistic basis for these alternative stable states in our model, we individually varied the dilution rate while allowing the pH to vary according to the accumulation of organic acids (see “Environment pH” section of the Supplementary Text S1). Two clearly distinguishable states emerged (Fig. 3A): one where *Blautia hydrogenotrophica* and *Bacteroides thetaiotaomicron* co-dominate, with *Roseburia intestinalis* almost completely outcompeted, and another where *Bacteroides thetaiotaomicron* and *Roseburia intestinalis* prevail, with *Blautia hydrogenotrophica* maintaining a low abundance.

**Fig. 3:**
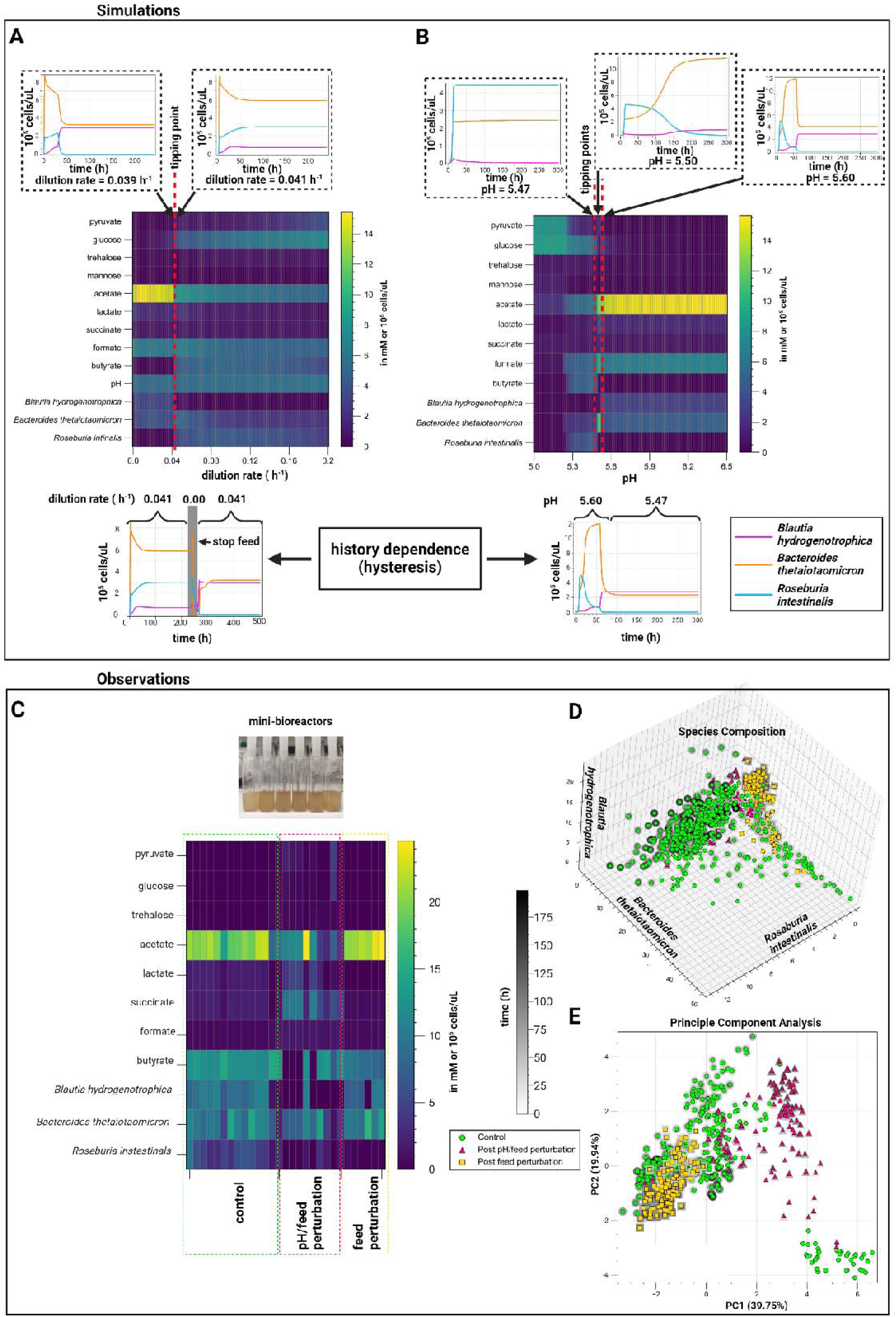
Simulations and experimental observations of tipping points and multistability in a gut microbial community. Model simulations predict alternative steady states connected by sharp transitions (tipping points) that exhibit hysteresis (ecological memory). For example, (**A**) a perturbation of the dilution rate drives the community towards an alternative state. Even when the original dilution rate is restored, the system does not return to its previous state. This scenario, depicted in the lower inset, was reproduced experimentally (see feed perturbation in panel C) (**B**) The steady-state community composition that emerges when controlling the system’s pH depends on the system’s history. For example, when the system starts and remains at a pH of 5.47, a different state is reached compared to when it starts at a pH of 5.60 and is shifted to a pH of 5.47. Notice that when varying the dilution rate (A), the pH is an emergent state of the system, whereas in the heatmap in (B), it is held constant at specific values. (**C**) Experimental validation conducted in minibioreactors (shown in the small inset photograph) reveals stable community compositions that persist under nearly identical environmental conditions, aligning with the model’s predictions of multistability. We systematically collected periodic samples and analyzed community states using HPLC for metabolites, alongside 16S rDNA sequencing and flow cytometry for bacterial species quantification in three independent experiments (detailed in Supplementary Table S2 and Figure 4). The heatmap illustrates the endpoint replicates of the control and perturbation setups. The 3D plot (**D**) shows the abundance of each species per time point, while the PCA plot (**E**) shows the principal components calculated from both the cell and metabolite concentrations for each replicate at each time point. The shadow around the points depicts time. We observe that following a feed perturbation, *Roseburia intestinalis* is outcompeted by *Blautia hydrogenotrophica* and *Bacteroides thetaiotaomicron*, while following sequential pH and feed perturbations, the system converges into different, unpredictable, states.

The shift to one state or the other depends on the concentration of *Blautia hydrogenotrophica*’s glucose-consuming cells. High dilution rates lead to trehalose accumulation, which in turn inhibits the glucose-consuming phenotype. When *Blautia hydrogenotrophica* is not consuming glucose, it occupies a niche that has negligible impact on the other species. In contrast, if the trehalose concentration is low and a sufficient population shifts to glucose consumption, *Blautia hydrogenotrophica* becomes a strong competitor, inhibiting the other species. Interestingly, this phenotype displays characteristics of hysteresis (ecological memory): once the glucose-consuming population is established, for example, by stopping the feed (setting the dilution rate to zero), then restoring the feed to its previous levels does not return the system to its former state (Fig. 3A). Consequently, two states can coexist under the same parameter values, and the system’s history is required to predict its behavior (for a more in-depth exploration of this behavior refer to Supplementary Fig. S3). We also observed this behavior experimentally (Fig. 3C, Fig. 4).

**Fig. 4.**
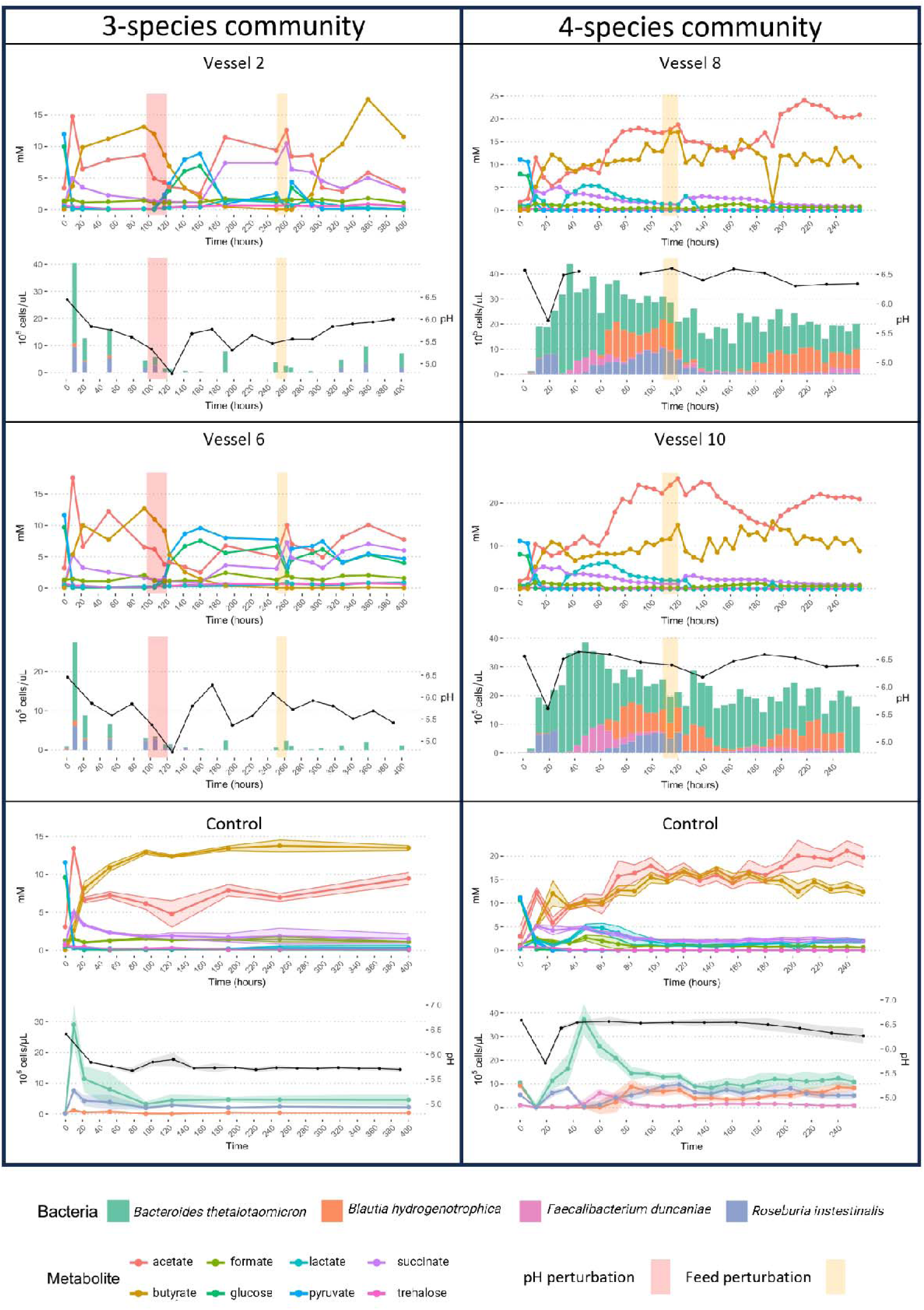
**Metabolite concentrations and bacterial abundances over time for two different experiments in minibioreactors**. A three-species community and a four-species community. Two vessels per perturbation experiment are shown, with the addition of control vessels (and standard deviation) for the same growth conditions without perturbations. The metabolite control for the three-species community consists of three replicates and the abundance and pH control plot consists of six replicates. For the four-species community, both controls consist of 12 replicates. Note that for the four-species experiment, *Segatella copri* was also inoculated but did not survive.

Similar alternative stable states also exist along the pH gradient. Because these states depend on the dilution rate, we fixed the dilution rate at 0.067 h^-1^ and explored the community landscape with varying pH values (Fig. 3B). At lower pH values (e.g. 5.47 see Fig. 3B), *Roseburia intestinalis* is favored. Overall, *Bacteroides thetaiotaomicron* is favored by pH control, since it can produce a large amount of organic acids without lowering the environmental pH and inhibiting its own growth. The state of the system, however, ultimately depends on its history, which is illustrated in the lower plot of Fig. 3B: changing the pH after the system is established does not necessarily lead to a change in community state (Fig. 3B). Notably, this effect is related to the concentration of cells in a specific phenotype reaching a tipping point and not to the extinction of *Roseburia intestinalis* cells, as the shift is performed while there is still a significant abundance of live *Roseburia intestinalis* cells in the system (also see Supplementary Fig. S3).

To test model predictions, we cultured gut species in minibioreactors that enable parallel cultivation and sampling in controlled conditions, emulating a chemostat via continuous inflow and pipette-controlled outflow, as reported previously ^27^. When a feed perturbation was applied, the system shifted towards a *Blautia hydrogenotrophica* and *Bacteroides thetaiotaomicron* enriched state as predicted by the model (yellow squares in Fig. 3D and 3E and Fig. 4B). These states show sharp transitions in species composition within a narrow range of experimental conditions, as we predicted with the model’s stability landscape (Fig. 2C). In reactors where *Blautia hydrogenotrophica* was highly abundant, trehalose was completely consumed (Fig. 4 and Supplementary Table 2), suggesting that the glucose-consuming subpopulation could emerge and influence the abundance of other species. Conversely, *Blautia hydrogenotrophica* was absent or present at low levels in reactors where trehalose remained at detectable concentrations.

In summary, perturbations have a lasting effect, even when the system is returned to its previous conditions. Following only a dilution rate perturbation (feed stop), the system transitioned to a state with high abundance of *Bacteroides thetaiotaomicron* and *Blautia hydrogenotrophica* and low abundance of *Roseburia intestinalis*, similar to the behavior predicted by the model (Fig. 3A and Fig. S3A). Following sequential pH and dilution rate perturbations—acidifying the pH to around 4.8 and later stopping the feed (Fig. 4A, Supplementary Fig. S4 and S6 and Supplementary table S2)—the replicates diverged into three alternative states after the system was returned to its previous conditions: either reverting to its previous state with low *Blautia hydrogenotrophica* abundance, transitioning to a state were the three species coexist, or moving to a state where *Bacteroides thetaiotaomicron* predominates. Since replicates from different experiments cluster into distinct steady states following perturbations (e.g. red triangles in Fig. 3E), this corroborates the history-dependent multistability mechanism suggested by our model’s landscape analysis.

## Simulating multistability in systems with many species

To further confirm this multistability mechanism and explore its potential to explain microbiome landscape dynamics, we abstracted key components of our model into a new formalism (for details, refer to Supplementary Text S2). Species are defined by Lotka-Volterra growth rates and interactions, but instead of single growth rates and interactions, species now encompass subpopulations with alternative phenotypes—two and potentially more growth rates and interaction coefficients—with environment-responsive transitions between them (Fig. 5A). As in our mechanistic model, this simplified model exhibits alternative community types separated by a tipping point (Fig. 5B). These states arise from subpopulation shifts to strongly competing phenotypes (e.g. more efficient usage of key nutrients). Simulations show that even in larger communities, environment-driven emergence of such competitive phenotypes can significantly reshape the community landscape, producing distinct community types that resemble enterotypes, i.e. alternative community types observed in fecal samples^3^ (Fig. 5C). Similar to an empirical study of species distribution across stool samples^28^, the species driving such community shifts—referred to by their original authors as “tipping elements”—exhibit a bimodal distribution in our model (Fig. 5C).

**Fig. 5:**
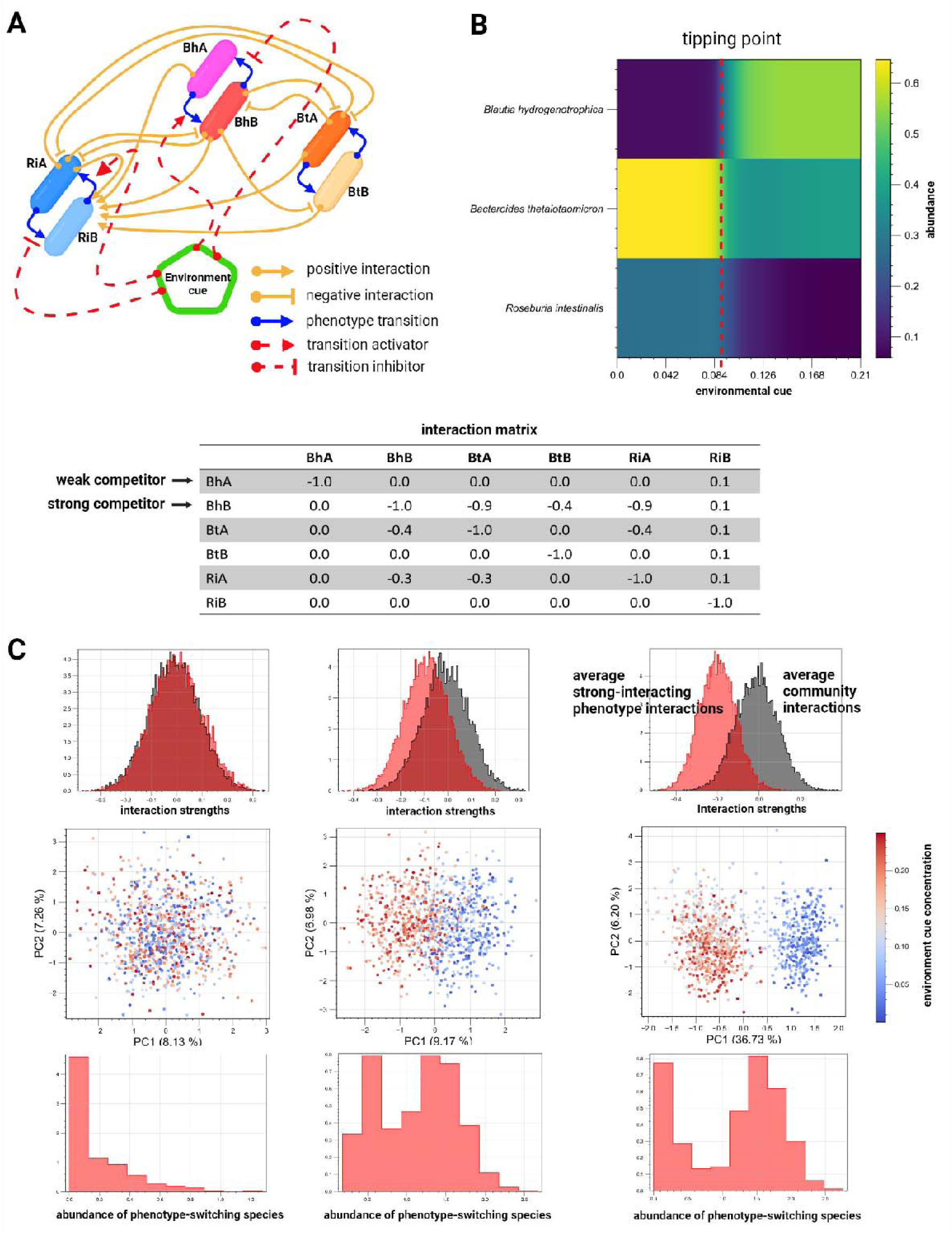
Toy model demonstrates conceptual mechanism for multistability. (**A**) Species interact through alternative phenotypes connected by environment-responsive transition functions, implemented through Hill equations, allowing dynamic switching between phenotypes during simulations (refer to Supplementary Text S2 for details). (**B**) If one phenotype strongly interacts with others (average interaction strengths are higher than the average community interactions), phenotype switching can induce a sharp transition between alternative community states (e.g. high steady-state trehalose leads *Blautia hydrogenotrophica* to a weakly competing phenotype, but low trehalose concentration triggers a metabolic shift, enabling *Blautia hydrogenotrophica* to strongly outcompete others in glucose). (**C**) Simulations with 1,000 random communities containing 50 species and random concentrations of an environmental factor show that this mechanism can explain emergent alternative stable states reminiscent of enterotypes (visualized as two distinct clusters in principal component space). Gray histograms show the distribution of interaction strengths across communities; red histograms show the distribution of interactions of a strongly interacting phenotype that is expressed in response to the concentration of an environmental factor. The lower histograms show the distribution of abundance of the species expressing the strongly interacting phenotype across samples. Notably, when enterotypes emerge, the switching species tends to exhibit a bimodal distribution across samples.

## Discussion

Here, we explored in depth the metabolic strategies of three prevalent human gut bacterial species and demonstrated in silico and in vitro that multistability is present and according to our model arises as a consequence of bacterial metabolic flexibility. Sharp transitions between alternative states in our system are driven by varying ecological interactions among phenotypically flexible bacteria. For example, we hypothesize that the glucose-consuming phenotype of *Blautia hydrogenotrophica* competes strongly with and inhibits the fast growth mode of *Roseburia intestinalis*. Communities with a low trehalose concentration—indicating that this phenotype is active—are significantly different from communities where it is repressed. The history-dependent behavior observed in our model and chemostat experiments emerges from feedback loops between subpopulations and environmental factors. For example, while increasing the dilution rate may increase the steady-state concentration of trehalose and inhibit *Blautia hydrogenotrophica*’s glucose-consuming phenotype, if a large population of glucose-consuming cells is already present in the system, then many cells are available to switch and quickly consume the excess trehalose that would otherwise accumulate due to the increase of the dilution rate. This leads to a different proportion of subpopulations and alters the community phenotype. Importantly, these changes have effects similar to those observed when the community is inoculated with vastly different species proportions^29^. However, in our system, these shifts are triggered by changes in environmental parameters and the associated microbial phenotypes. This mechanism is likely closer to real-world conditions but involves subtle environmental changes that can be difficult to predict. A limitation of our work is that we did not collect direct evidence of phenotype switching in the community, which requires stable isotope experiments and is a task for the future.

Our simulations further confirm that the presence of different phenotypes can give rise to alternative community types in large communities. Thus, we suggest that multistability is a potential driver behind alternative community types observed in the gut microbiome^1,3^, supporting previous propositions^1,7^. The occurrence of alternative community types in other host-associated microbiota, which are not easily explained by environmental differences^2,30,31^, implies that multistability may be more common than previously thought. As we and others have previously discussed^7,8,28^, it is relevant whether alternative community types are due to environmental differences caused by diet, host genetics, or other factors or whether they are due to multistability. In the case of the latter, alternative community types can result from past transient perturbations rather than from current differences between hosts.

We also note that popular mathematical models of microbial communities, such as the generalized Lotka-Volterra (gLV) model, do not account for metabolic flexibility. Moreover, several established methods assume the absence of multistability in microbial communities. One of these is the dissimilarity-overlap curve analysis^32,33^, which evaluates the universality of microbial interactions by relying on an empirical negative correlation between compositional dissimilarity and species overlap. Another is EPICS (effective pairwise interactions for predicting community structures), which parameterizes the gLV model from leave-one-out communities^34^. The accuracy of other inference methods such as BEEM^35^ or MDSINE^36^ may also be affected by the occurrence of multistability.

There are different ways to integrate metabolic knowledge into community models^37^. Here, we opted for a kinetic model instead of a metabolic model. This choice was made to effectively capture pH response and phenotypic switches, as well as to investigate history-dependence and the stability landscape of the community. However, the kinetic model was designed based on insights manually derived from metabolic reconstructions. It may be possible to construct such kinetic models automatically from metabolic models in the future.

In summary, we have shown that flexible microbial strategies impact the composition of gut microbial communities. In the future, we need to systematically elucidate these strategies in other gut microorganisms^38^ to better understand and efficiently modulate gut microbial communities.

## Acknowledgements

We thank Gertjan Gerits for helping in the experiments shown in Fig. S1B and Aristeidis Litos for some useful discussions regarding the model in Fig 4A. We would like to thank Emma Hernandez-Sanabria, Veronica Lloréns-Rico and Jelle Matthijnssens for input and support concerning bacterial transcriptomics, as well as Jeroen Raes for access to the CytoFLEX. We are also grateful to Thi Thuy Duyen Nguyen, Leen Rymenans and Geert Huys for assistance in flow cytometry, 16S rRNA gene library preparation and anaerobic cultivation, as well as to Raul Yhossef Tito Tadeo for processing raw sequence reads.

## Author contributions

Conceptualization: KF, DRG, CV, DG Data curation: BL, CV, DRG, KS

Formal analysis: DRG, BL, KS, CV

Funding acquisition: KF, KB Investigation: BL, CV, DRG, PS, XZ Methodology: all authors

Project Administration: KF Resources: KF, KB, KS Software: DRG Supervision: KF Validation: all authors Visualization: DRG, BL

Writing—original draft: DRG, KF, BL Writing—review & editing: all authors

## Data and materials availability

All data the data and code used in the analysis is available at the GitHub repository: https://github.com/danielriosgarza/hungerGamesModel, which include detailed instructions to reproduce all the Figures in the manuscripts (https://github.com/danielriosgarza/hungerGamesModel/blob/main/multistabilitymanuscript.i pynb).

The raw RNA-seq data was deposited in the Sequence Read Archive (SAMN39333017-19 - https://www.ncbi.nlm.nih.gov/bioproject/PRJNA1063153/; SAMN32321133-38, https://www.ncbi.nlm.nih.gov/bioproject/PRJNA914119/).

Raw flow cytometry data is deposited in flowrepository.org (IDs FR-FCM-Z6YM, FR-FCM- Z6YN, FR-FCM-Z74P, FR-FCM-Z753 and FR-FCM-Z754).

**Supplementary Materials** Materials and Methods Supplementary Text S1^24,26,39–41^ Supplementary Text S2

Supplementary Figures S1, S2, and S3 Supplementary Movie S1

## Online Methods

### Microbial strains

Human gut bacterial strains of *Blautia hydrogenotrophica* S5a33 (DSM 10507^T^), *Bacteroides thetaiotaomicron* VPI-5482 (DSM 2079^T^) and *Roseburia intestinalis* L1-82 (DSM 14610^T^) were obtained from the Deutsche Sammlung von Mikroorganismen und Zellkulturen (DSMZ, Germany). The strains were frozen in Wilkins-Chalgren Anaerobe Broth (WC; Oxoid Ltd., Basingstoke, United Kingdom) plus 20% glycerol and maintained at -80°C until use.

### Batch cultivation and sample collection

Batch cultivations of monoculture, bi-culture and tri-culture were followed for 120 h in 120- ml serum bottles containing 60 ml of WC medium. The serum bottles were prepared in a same way as previously described^24^, and were inoculated with 1 ml of the diluted preculture to an OD600 of 0.1 (either a single species or the mixture of them). The bottles were incubated at 37°C and at a constant stirring rate of 170 rpm (shaker KS 4000 i; IKA, Staufen, Germany).

Samples were taken from the liquid broth every four hours for the first 48 h and every 12 h afterwards. Three biological replicates were designed for testing the monocultures in three independent batch experiments. All bi-culture and tri-culture experiments were performed in six biological replicates and always had a negative control bottle without inoculation but with sampling for each time, to verify its sterility.

Sterile syringes were used for each timepoint to collect 1 ml of the fermentation broth into 2- ml tubes (Eppendorf) under anoxic conditions. Subsequently, these tubes were used to measure OD600, pH, metabolites and to count bacterial cells by live/dead staining followed by flow cytometry.

### Chemostat experiment with ambr 15

Fermentations were performed in the Ambr® 15 Fermentation system (Sartorius Stedim Biotech, Royston, UK) located inside a Don Whitley A155 Anaerobic Workstation with HEPA filter (10% H_2_, 10% CO_2_, 80% N_2_, 55% humidity) as previously described^27^. Strains reported were precultured first for 48 h, in modified Gifu Anaerobic Medium broth (mGAM, HyServe) then cultured for 18 hours in WC (1/100th dilution without washing) before inoculating the minibioreactors.

Prior to inoculation, strains were diluted in WC medium, mixed in even ratios based on OD_600_ and inoculated in the minibioreactors to a total volume of 10 ml (OD 0.001 for *Bacteroides thetaiotaomicron* and 0.002 for the others). A sample was taken at time point 0 and continuous feeding and sampling started at 4 hours after inoculation. The feed consisted of WC anaerobe broth and was delivered at an approximate rate of 0.04 h^-1^, resulting in a complete change of the medium in 24 hours. Samples (250 μl) were pipetted into a cooled plate (4°C).

In the first two of the three experiments (Supplementary table S2) *Segatella copri* (DSM 18205, previously Prevotella) and *Faecalibacterium duncaniae* (DSM 17677, previously *F. prausnitzii*) were also inoculated into the system. However, they maintained negligible abundance. We report absolute abundances based on flow cytometry measurements and confirmed the main observation of multistability following a sequential pH and feed perturbation inoculating the system with only *Blautia hydrogenotrophica*, *Bacteroides thetaiotaomicron*, and *Roseburia intestinalis*.

In the first experiment, we first applied a pH perturbation by decreasing the pH of the feed (WC with pH 6.4 to pH 3.7) between 88 and 112 hours (24 hours total). Subsequently, we applied a perturbation in the dilution rate by stopping the feed for 12 hours between 150 and 162 hours (periodic removal of liquid continued), after which an additional 5 mL of fresh medium (50% of total volume) was added to the vessels.

In the second experiment, we applied only a feed perturbation between 108 and 120 hours (12 hours) as described above, including the 5mL addition of fresh medium (Supplementary Fig. S6).

In the third experiment, we applied a pH perturbation by decreasing the pH of the feed (WC with pH 6.4 to pH 3.38) between 97 and 109 h (24 hours total). Subsequently, a feed perturbation was applied between t251 and t263 (12 hours) as described above including the addition of fresh medium.

The 16S rRNA genes of selected samples was extracted and sequenced as previously described^27^. Additionally, we used Nanopore MinION flow cell sequencing in our last minibioreactor experiment (Supplementary Fig. S6). For this purpose, genomic DNA was extracted using ZymoBiomics DNA miniprep kit (Zymo, USA) with slight modifications to prevent biased extraction. After DNA extraction, the full-length 16S rRNA genes were amplified through a single PCR program. Zymo Biomics Microbial Community DNA standard (Zymo, USA) is included in one well per sequencing plate as the quality control to avoid any bias in different sequence libraries. The rest of operations were carried out according to the protocol of EasySeq™ Full-length 16S Library Prep Kit (NimaGen, Netherlands). The sequence library was then loaded on Nanopore MinION flow cell and sequenced by MinKNOW (version 24.02.8). The minimal read-length was set to 1,000 bp and real-time basecalling was operated with the super-high accuracy model to improve sequence quality. The barcoding score was set to at least 70 to demultiplex and trim the barcode afterward. The long-read classifier minimap2 was employed with the NCBI RefSeq 16S rRNA database for taxonomic classification^42,43^. The 16S rRNA gene copy number database rrnDB is used to correct the different 16s rRNA copy numbers of the strains in the community^44^.

### Quantification of live cells with flow cytometry

We used a combination of the DNA-based stains SYBR Green I (SG; Invitrogen) and propidium iodide (PI; Invitrogen) to stain bacterial cells with intact and damaged cytoplasmic membranes^45^. Under anoxic conditions, cells were diluted in filter-sterilized PBS buffer. 1:10 for the first two time points (0 and 4 h) and 1:200 for the next points and stained with a saturating solution of SG/PI, incubated for 20 min in the dark at 37 °C right and immediately measured by flow cytometry using the benchtop CytoFLEX S flow cytometer (Beckman Coulter, Brea, USA) instrument. Events were recorded for exactly 1 min at a sample flow rate of 10 μl/min, applying threshold values of 3000 and 2000 for the forward and side scatters, respectively, values that we have previously validated^24^. We also used 0.5 μm and 1 μm green fluorescent beads (Thermo Fisher Scientific, USA) as internal standards.

### Flow cytometry data analysis

We used an in-house developed pipeline to accurately quantify the absolute abundance of live *Blautia hydrogenotrophica*, *Bacteroides thetaiotaomicron*, and *Roseburia intestinalis* cells for batch cultures. We clustered flow cytometry events in a UMAP space, following a detailed protocol available at: bit.ly/3WNrslL. Raw flow cytometry data were normalized, scaled, and transformed using the arcsin function. The data was then projected into three-dimensional UMAP space, allowing us to classify cell populations into four categories: “live,” “inviable,” “debris,” and “blank.” This classification was based on empirically determined thresholds for propidium iodide (PI) and SYBR green (SG) signals and by distinguishing from blank control runs.

Further analysis involved using monoculture samples for species classification in co-culture samples. We created a training space with labeled UMAP projections of random events from each monoculture replicate. Supervised UMAP and the *K*-nearest neighbors vote classifier (parameters: n_neighbors = 50, weights = distance, and metric = Mahalanobis) were employed to assign species labels to co-culture events. Prior to finalizing classifications, each sample was overlaid with corresponding blank runs for manual verification using clickable three-dimensional scatter plots, ensuring accurate separation of cell populations from blanks. This process also validated our parameter choices for UMAP and the classifier. Additional details and scatter plot examples are available at: bit.ly/3WNrslL.

### Metabolite profiling

After centrifuging the liquid broth for 20 min at 21,130 × *g* at 4°C (Centrifuge 5424R; Eppendorf, Hamburg, Germany), the supernatants were used for measuring the concentrations of trehalose, glucose, pyruvate, succinate, formate, acetate, lactate and *n*-butyrate, which were determined by high-performance liquid chromatography (HPLC) as previously reported^24^. We also measured the first and end time points of the blank controls. Metabolites propionate, *iso*- butyrate, *n*-valerate and *iso*-valerate were measured but their concentrations were not consistently different from the blank WC control. Most of these metabolites are found in our system as organic acids. In the text, however, for brevity we refer to them by their salt names.

### RNA extraction and sequencing

A total of 27 samples representing the different growth phases of monocultures in three biological replicates were selected for RNA sequencing, including timepoints of three independent experiments: monoculture *Blautia hydrogenotrophica* in WC (14h, 32h and 72h), monoculture *Bacteroides thetaiotaomicron* in WC (4 h, 12 h and 36 h) and *Roseburia intestinalis* in WC (4 h, 12 h and 48 h). Details of the extraction and purification of total RNA, evaluation of RNA integrity and yield, as well as library preparation and sequencing can be found in our recent paper^24^. Although we used the same methodology, the RNA sequencing data of *Blautia hydrogenotrophica* is unique to the current study.

### RNA-seq data processing

Initially, low-quality reads and adapters were removed using fastp. We then mapped high-quality RNA reads to reference transcripts using Salmon in selective alignment mode, employing a decoy-aware index constructed from each organism’s genome. We used the latest reference genomes and transcripts from the BV-BRC database. from the counts files we used the R package DESeq2 (https://bioconductor.org/packages/release/bioc/html/DESeq2.html) to estimate the *p-*values between pairs of conditions. The R scripts for these analyses can be found here: https://github.com/danielriosgarza/hungerGamesModel/blob/main/scripts/R/

To integrate the gene expression data with genome annotations and metabolic information from the genome-scale metabolic models, we wrote a Python class, which is available here: https://github.com/danielriosgarza/hungerGamesModel/blob/main/scripts/geneExpression/par seGenExpData.py. With this class one can, for example, draw the bar charts of Fig. 1A, B, and C.

### Modeling

In this manuscript we built two computational models based on ordinary differential equations: a mechanistic model based on metabolite and cells kinetic equations (depicted in Fig. 2A) and a phenomenological model based on the generalized Lotka-Volterra dynamics (depicted in Fig. 5A). A detailed description of the model parameters, rationale, and experimental validation is available in the Supplementary Texts S1 and S2. A detailed implementation of the reported simulation and code to reproduce all of our computation analysis is available at the projects’s Github repository: https://github.com/danielriosgarza/hungerGamesModel, which also contains a comprehensive Wiki to help users reproduce our analysis, and detailed instructions to reproduce all the manuscript Figures in a Jupyter notebook (https://github.com/danielriosgarza/hungerGamesModel/blob/main/multistabilitymanuscript.i pynb).

---

**Supplementary Fig. S1:**
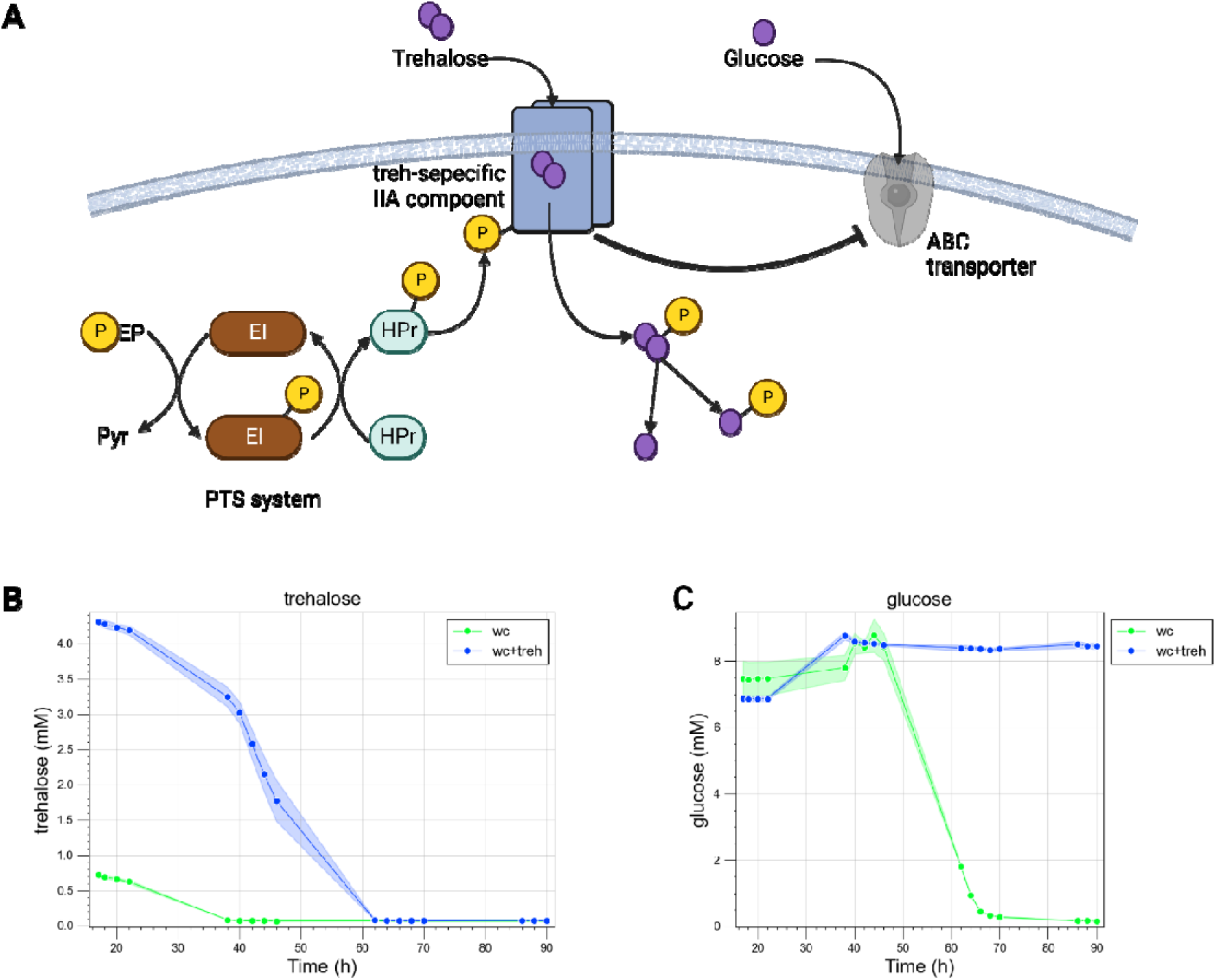
*Blautia hydrogenotrophica* neglects glucose when trehalose is present in the medium. (**A**) Illustration of a hypothetical PTS system that detects trehalose and inhibits the expression of the ABC transporter responsible for importing glucose. Although the detailed molecular mechanisms of this inhibition have not been fully elucidated, RNA-seq data clearly show inhibition of the ABC transporter when trehalose is present (refer to main Fig. 1A). Moreover, (**B**) supplementing WC medium with additional trehalose results in the (**C**) complete inhibition of glucose consumption (as compared with the ’glucose’ plot in main Fig. 1D).

**Supplementary Fig. S2:**
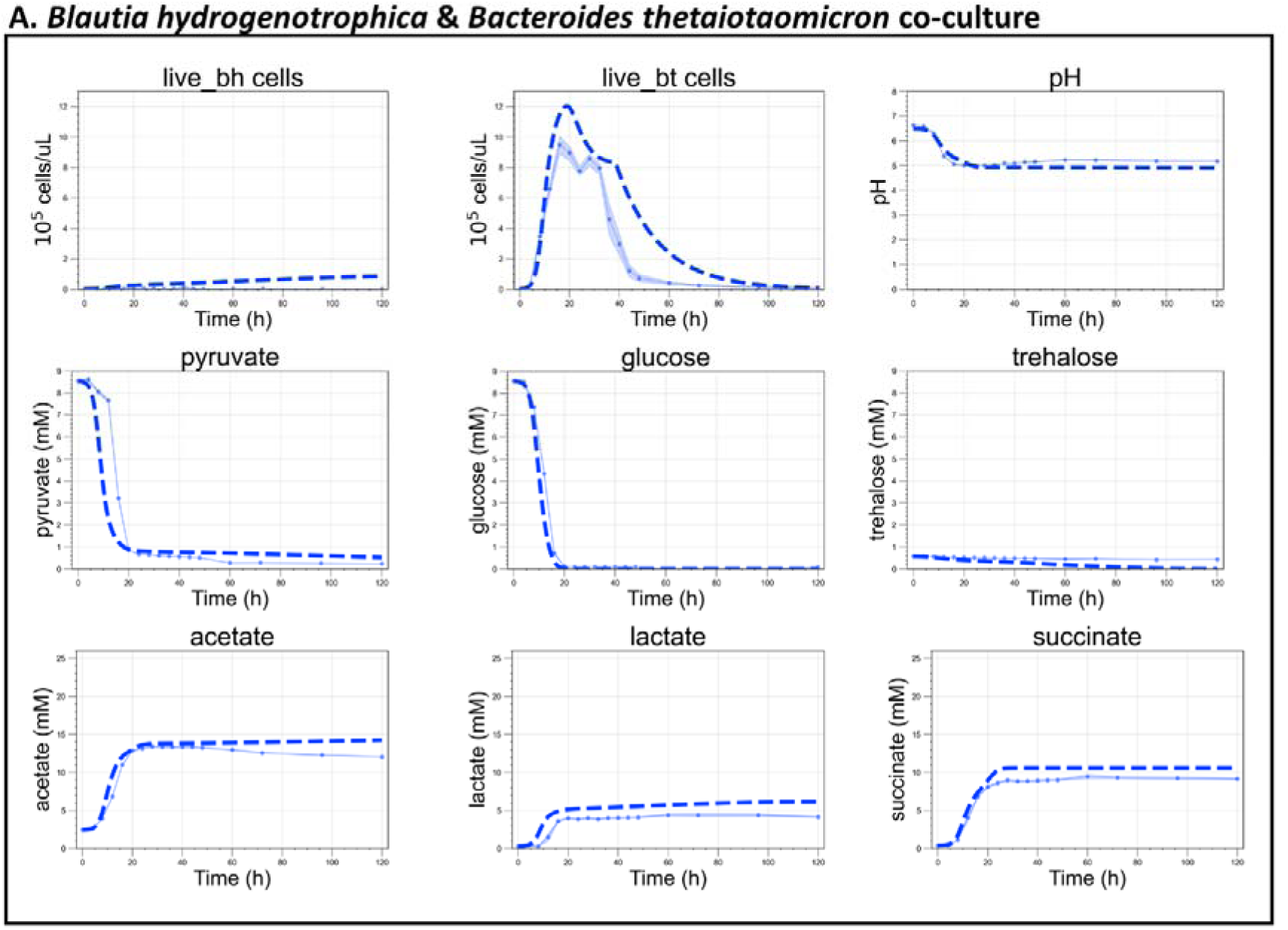
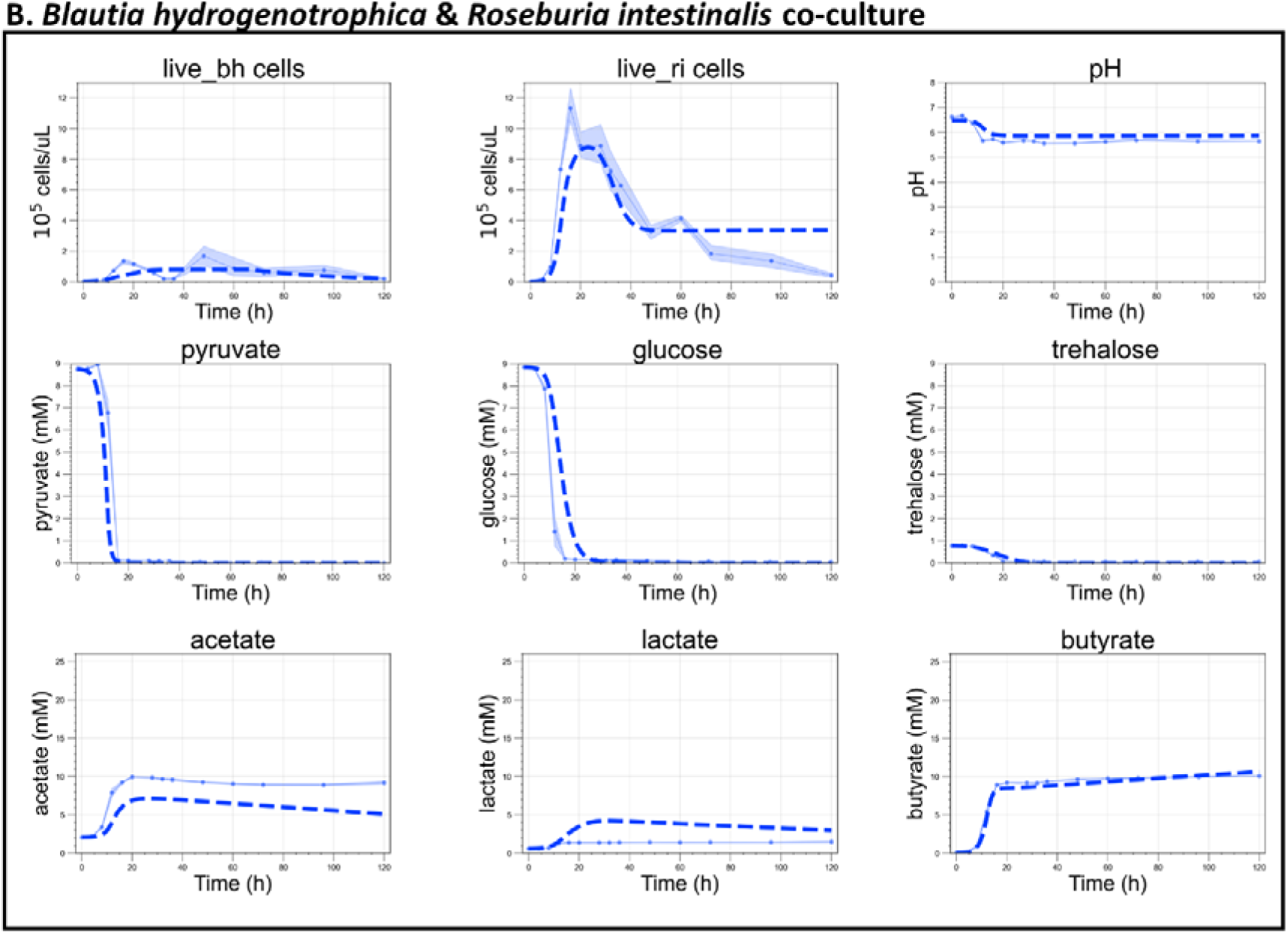
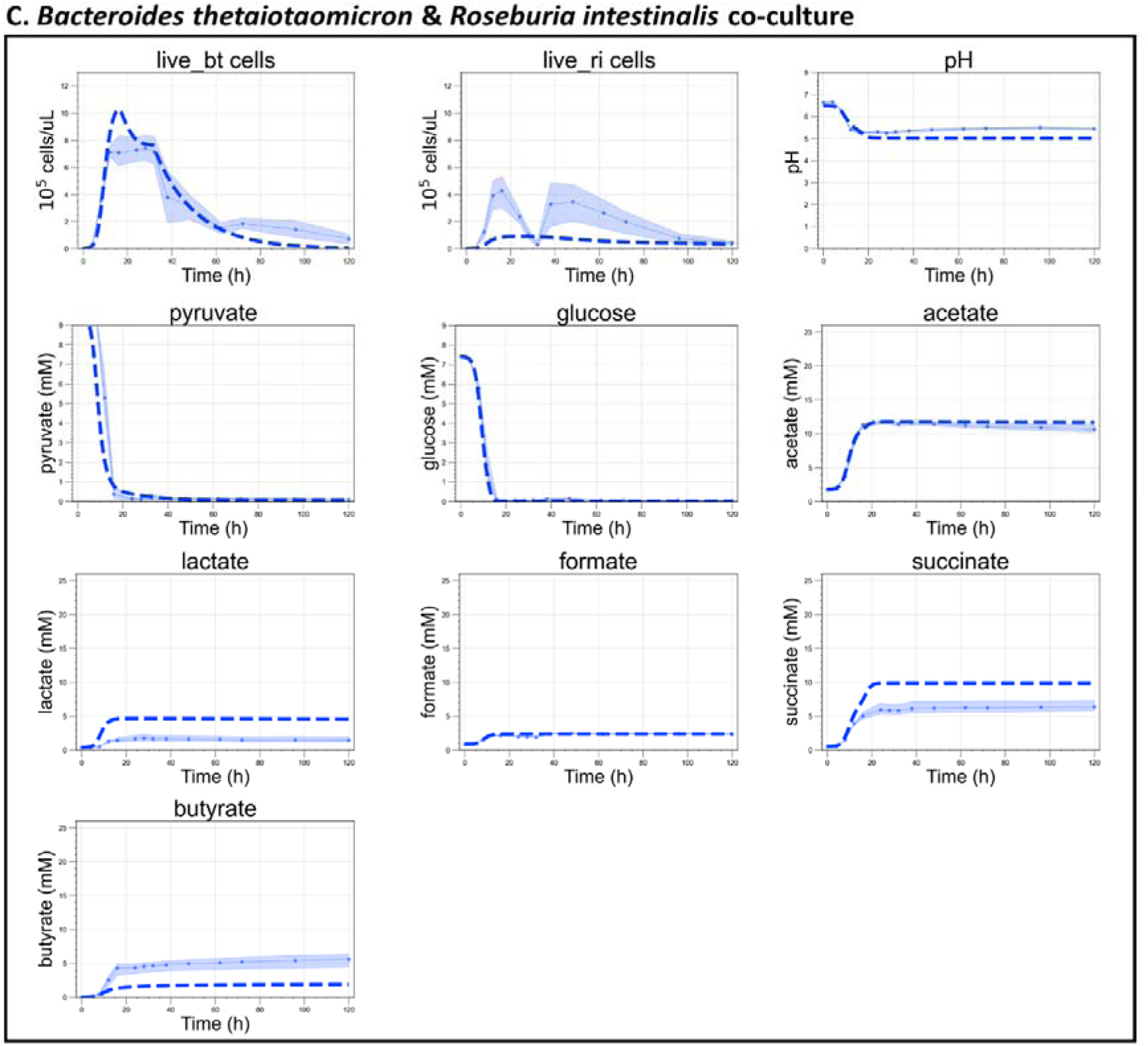
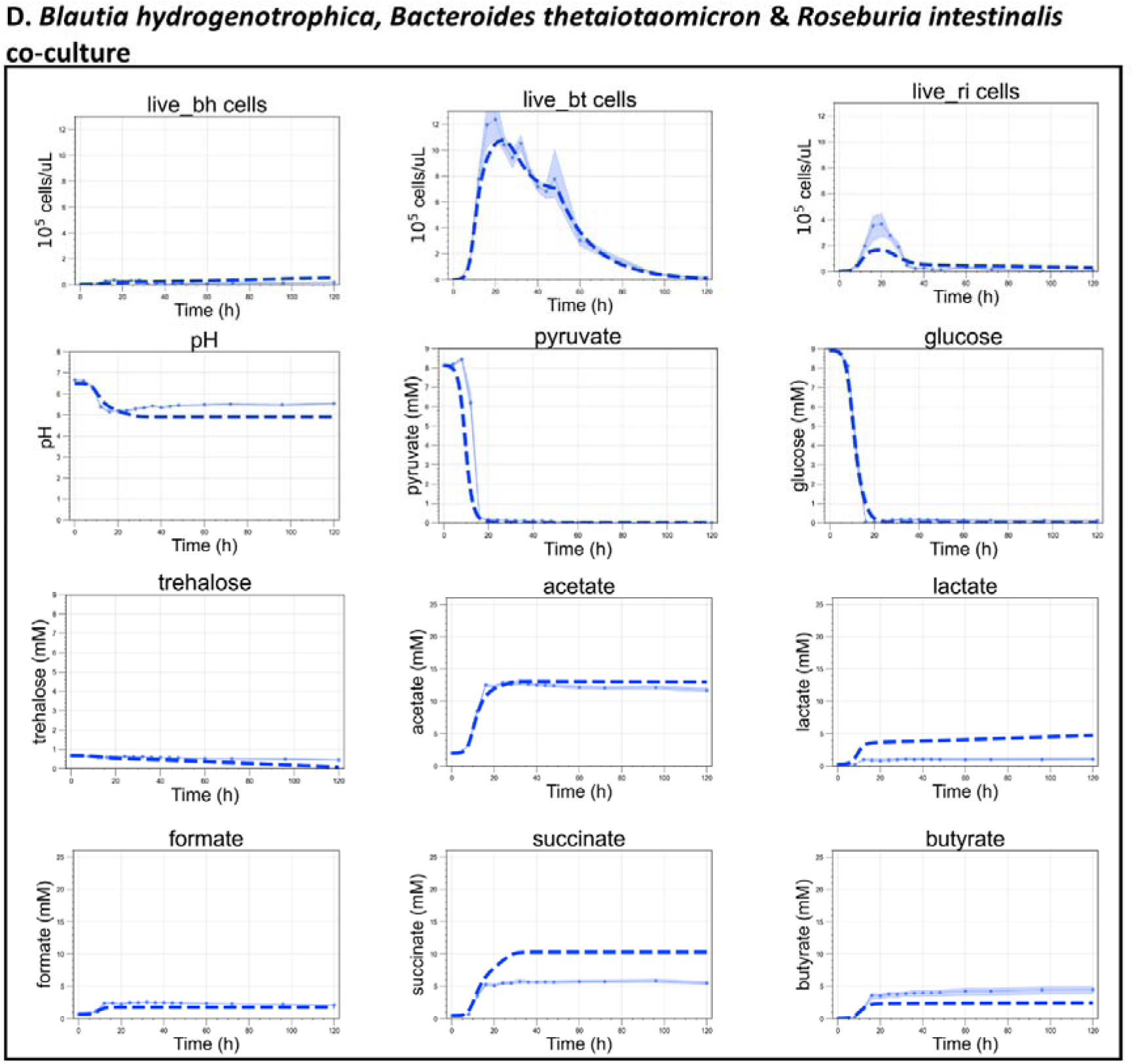
Growth kinetics and model validation for three human gut bacteria in co-culture. This figure presents experimental growth data over time alongside model simulations (indicated by dashed lines) for pairs of species and all three species co-cultured in WC medium. The growth data represent averages from six biological replicates. Simulation of initial conditions matched those of the experimental setups. In experiment (**A**), the HPLC analysis for formate failed, resulting in the absence of experimental data points. Only the metabolites that showed changes relative to the blank control are depicted. Other metabolites, such as propionate in all experiments, and trehalose in the co-culture of *Bacteroides thetaiotaomicron* and *Roseburia intestinalis*, were measured but remained at levels equivalent to the blank, which consisted of pure WC medium; hence, they are not included in these plots.

**Supplementary Fig. S3:**
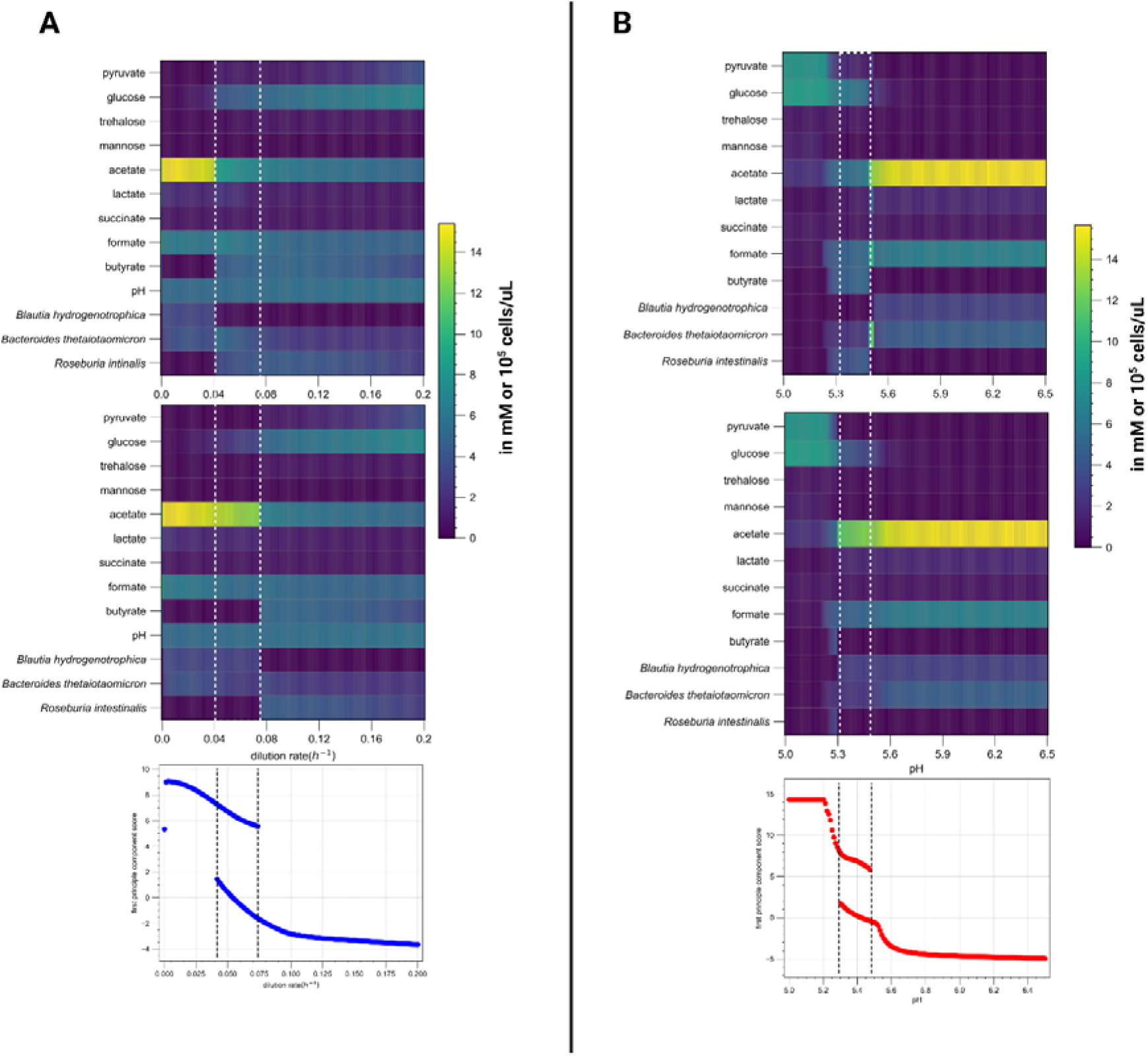
Two examples of history dependence in simulations using our mechanistic model. In (**A**), simulations began with dilution rates corresponding to the values on the x-axis and reached a steady state depicted in the top heatmap. After reaching this steady state, the dilution rate was set to zero for 24 hours and then restored to its previous value, allowing the system to achieve a second steady state up until 600 hours. This second steady-state is shown in the lower heatmap. Within a certain range of dilution rate values (indicated by the area enclosed in the white traced lines), the system settles into a different steady state than in the case where no perturbation was introduced (notice, the higher concentration of *Blautia hydrogenotrophica* and acetate and lower concentration of *Roseburia intestinalis* and butyrate). The two steady states have the same parameter values and emerge depending on the history of the system. This alternative steady state is maintained by feedback loops among phenotypically distinct subpopulations. For instance, if the initial population of *Blautia hydrogenotrophica*’s glucose-consuming cells exceeds a tipping point, an increase in the steady-state concentration of trehalose (achieved by increasing the dilution rate) prompts *B. hydrogenotrophica* cells to switch to consuming the excess trehalose, keeping trehalose levels low and allowing glucose-consuming cells to persist. In other words, whether glucose-consuming cells persist or not depends on their initial population, which in turn depends on the history of the system. Using principal component analysis (PCA), the model state can be summarized by a single state value (its first principal component). In the lower plot, one clearly sees the coexistence of two states under the same parameter regime within the traced line (top unperturbed and bottom perturbed states). (B) A similar example of history dependency can be seen by controlling the system’s pH at specific values (for instance, by adding a buffer). In these simulations, in the top heatmap we show the steady state reached by the system when the pH is fixed from the beginning at the values in the x-axis. While in the lower one, we instead initialize the system with a fixed pH of 5.6 for 60 h. A short time period that would not lead *Roseburia intestinalis* to extinction and not necessarily enough time to reach a steady state. After 60 h, we set the pH to the specific value in the x-axis and allow the system to reach steady state (for 600 h). Here, once more within the range indicated by the white traced lines, two alternative states were reached that depend on the system’s history. In other words, the concentration of subpopulations that emerge after the initial incubation at pH 5.6 is more robust to pH changes and stabilizes the system in a different state compared to the initial populations of a standard incubation.

**Supplementary Figure S4.**
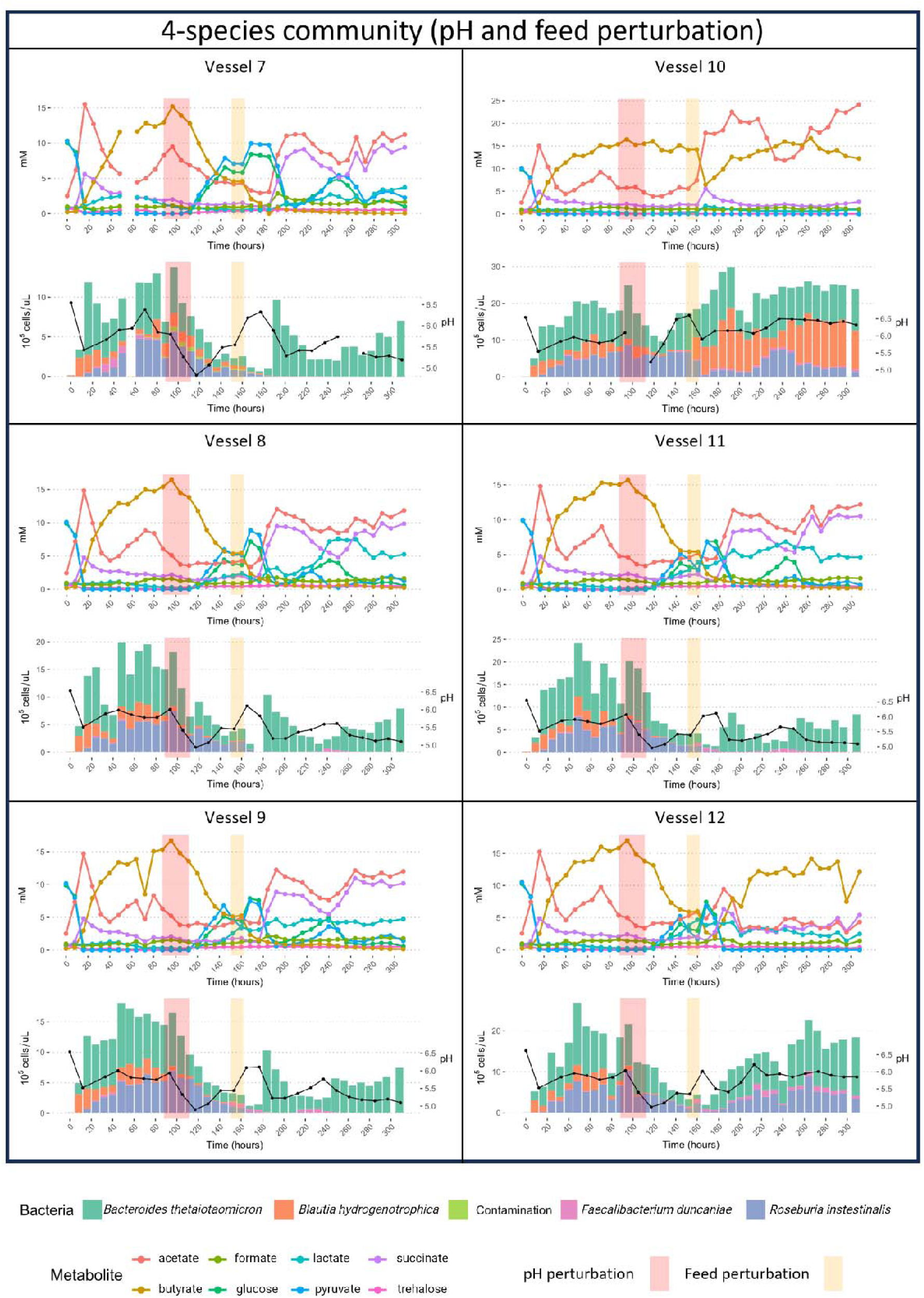
: Metabolite concentrations, bacterial abundances and pH for a 4- species community grown in minibioreactors. Relative abundances from 16S rRNA gene sequencing (copy-number corrected) were multiplied by total cell count from flow cytometry to obtain absolute abundance for six replicate vessels. Cells were grown in batch for 4 hours after which the emulated chemostat conditions were initiated. The red squares indicate the start and end of the pH perturbation, and the orange squares indicate the start and end of the starvation period (no feed added) followed by the addition of 5 ml medium.

**Supplementary Figure S5:**
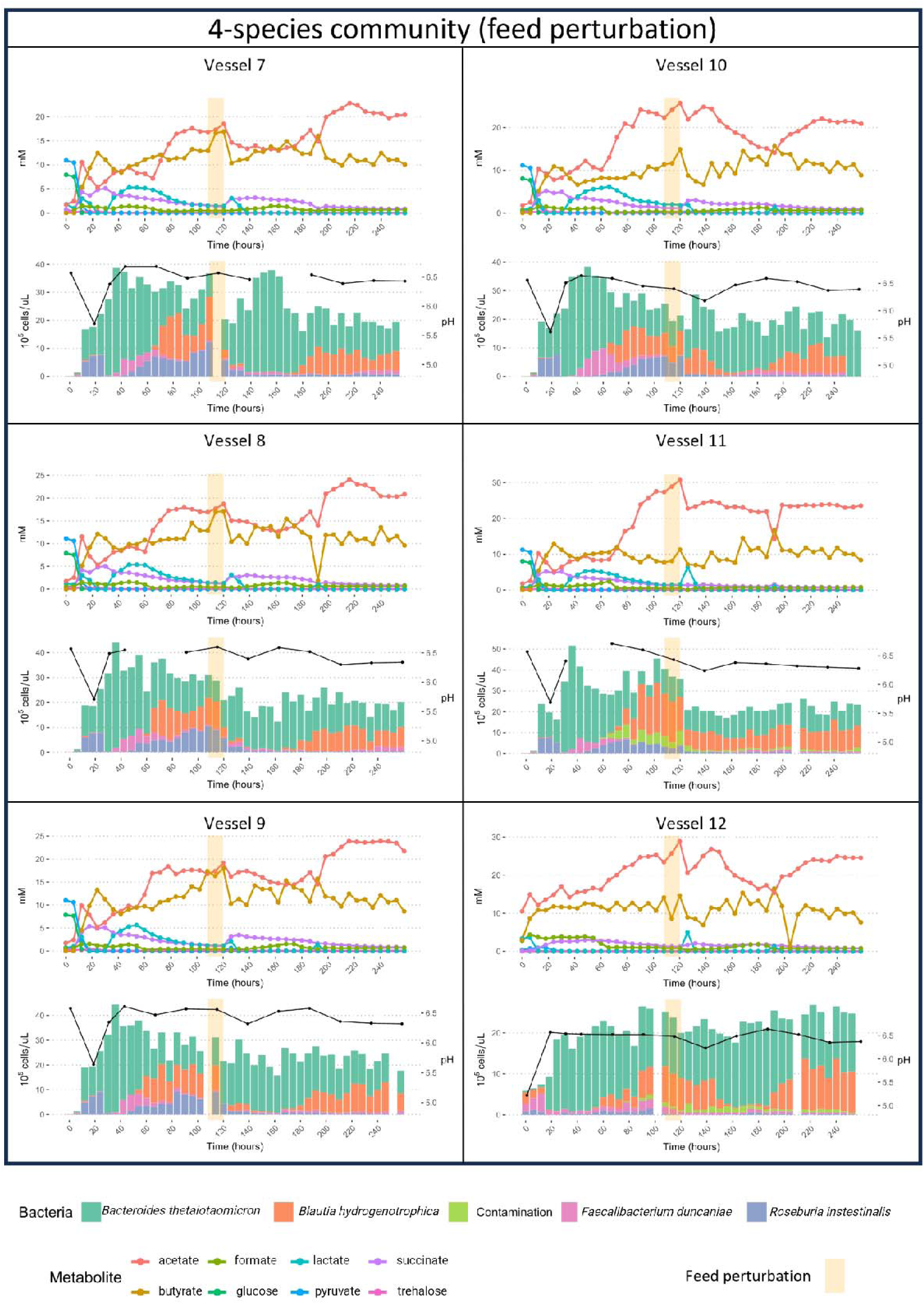
Metabolite concentrations, bacterial abundances and pH for a 4- species community grown in minibioreactors. Relative abundances from 16S rRNA gene sequencing (copy-number corrected) were multiplied by total cell count from flow cytometry to obtain absolute abundance for six replicate vessels. Cells were grown in batch for 4 hours after which the emulated chemostat conditions were initiated. The orange squares indicate the start and end of the starvation period (no feed added) followed by the addition of 5 ml medium.

**Supplementary Figure S6.**
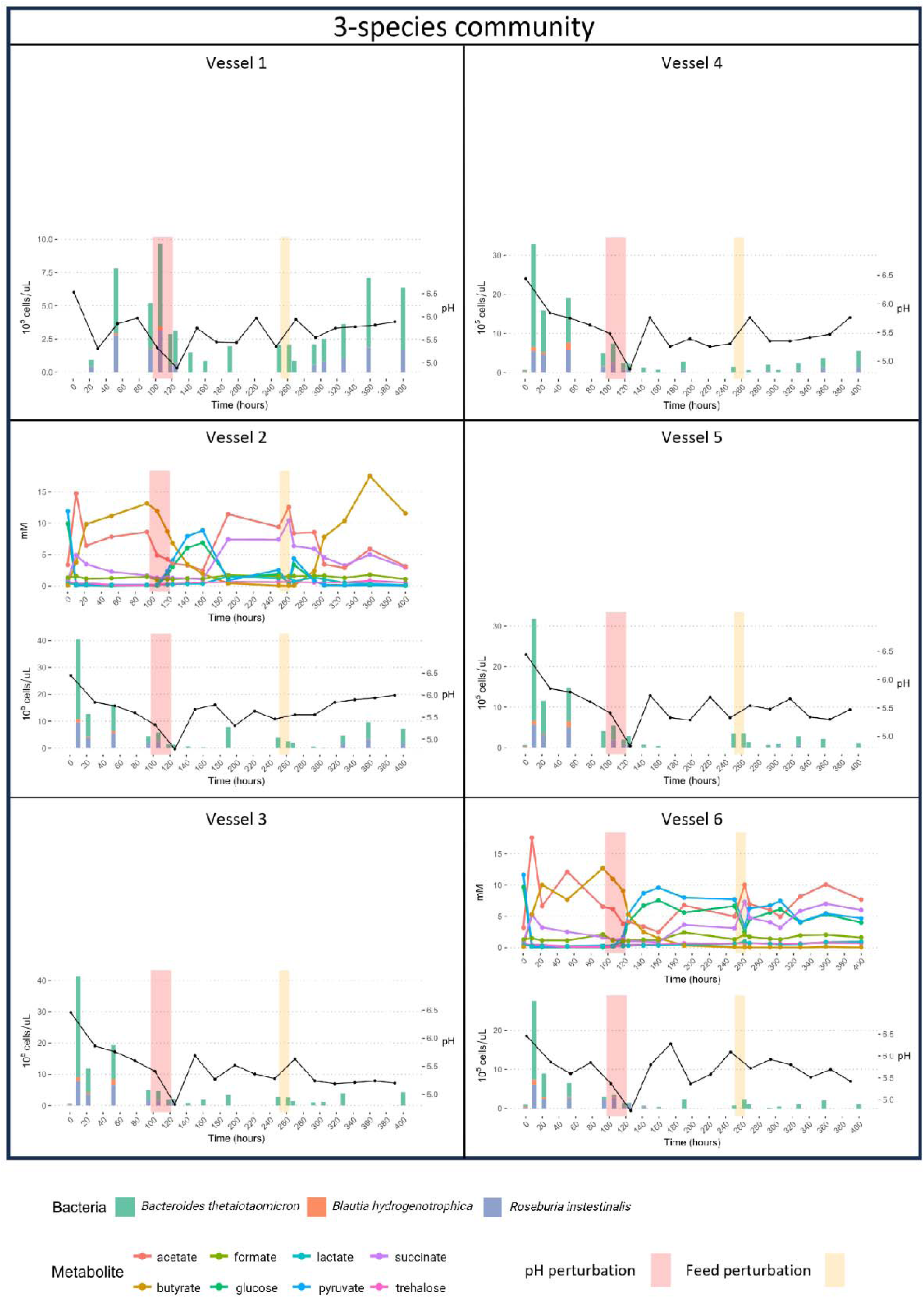
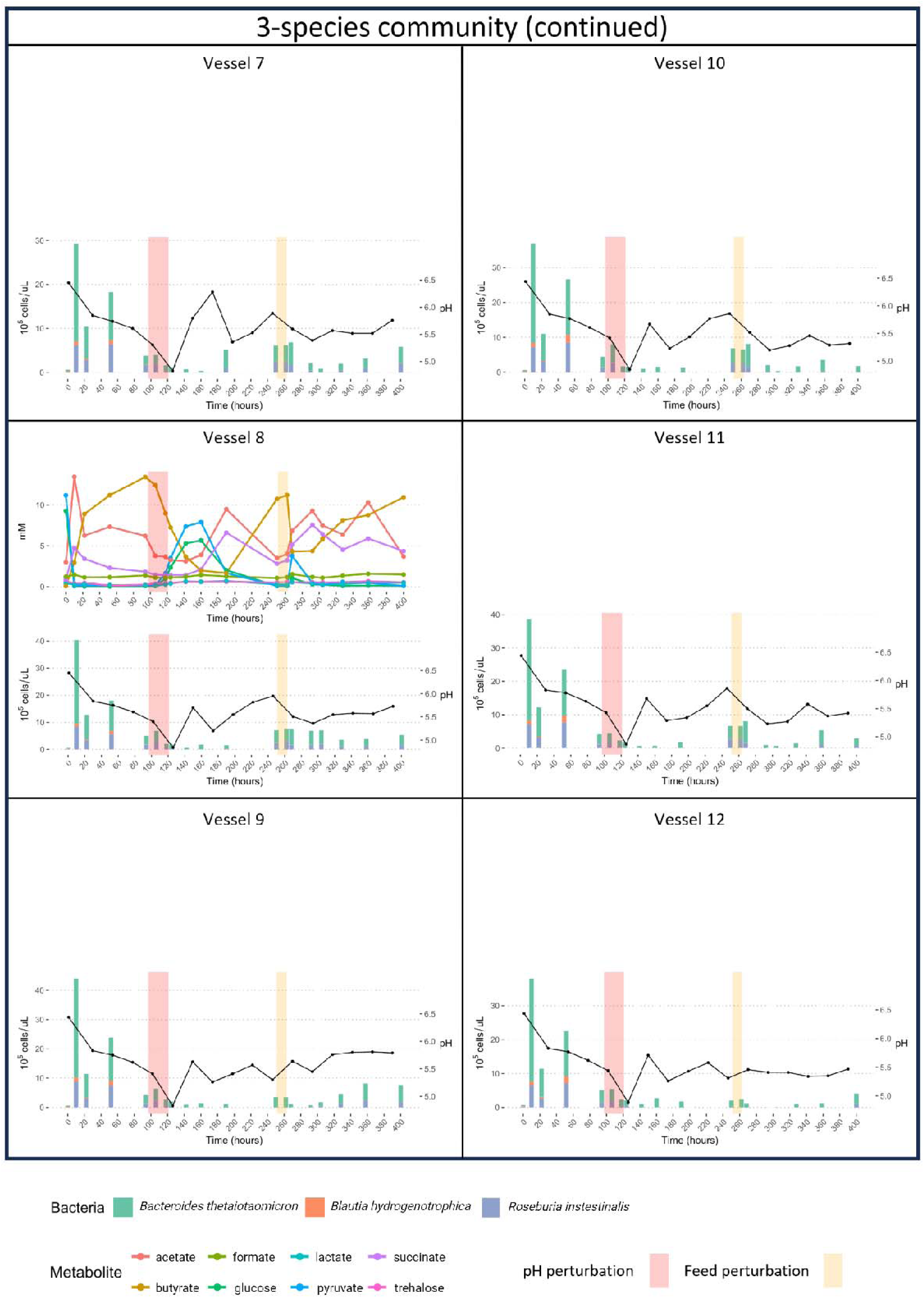
: Metabolite concentrations, bacterial abundances and pH for a 3- species community grown in minibioreactors. Relative abundances from 16S rRNA gene sequencing (copy-number corrected) were multiplied by total cell count from flow cytometry to obtain absolute abundance for six replicate vessels. Cells were grown in batch for 4 hours after which the emulated chemostat conditions were initiated. The red squares indicate the start and end of the pH perturbation, and the orange squares indicate the start and end of the starvation period (no feed added) followed by the addition of 5 ml medium.

**Supplementary Movie S1**: flow cytometry time series of *Blautia hydrogenotrophica*’s growth in WC medium, corresponding to experiment 1 shown in main Fig. 1D. ‘t’ represents the hours of cultivation. The x-axis displays the intensity of the SYBR green staining, indicative of viable (non-permeable) cells, while the y-axis shows the intensity of propidium iodide staining, which marks non-viable (permeable) cells. Between 24 and 32 hours, two distinct subpopulations of similar sizes become apparent, which coincides with the uptake of glucose (see main Fig.1D). These subpopulations were not observed in the experiments where WC medium was supplemented with trehalose (see Supplementary Fig. S1).

## References

1. Arumugam, M. et al. Enterotypes of the human gut microbiome. Nature 473, 174–180 (2011).

2. Gajer, P. et al. Temporal Dynamics of the Human Vaginal Microbiota. Science Translational Medicine 4, 132ra52-132ra52 (2012).

3. Costea, P. I. et al. Enterotypes in the landscape of gut microbial community composition. Nat Microbiol 3, 8–16 (2018).

4. David, L. A. et al. Host lifestyle affects human microbiota on daily timescales. Genome Biology 15, R89 (2014).

5. Dethlefsen, L. & Relman, D. A. Incomplete recovery and individualized responses of the human distal gut microbiota to repeated antibiotic perturbation. Proceedings of the National Academy of Sciences 108, 4554–4561 (2011).

6. Gibson, T. E., Bashan, A., Cao, H.-T., Weiss, S. T. & Liu, Y.-Y. On the Origins and Control of Community Types in the Human Microbiome. PLOS Computational Biology 12, e1004688 (2016).

7. Gonze, D., Lahti, L., Raes, J. & Faust, K. Multi-stability and the origin of microbial community types. ISME J 11, 2159–2166 (2017).

8. Goyal, A., Dubinkina, V. & Maslov, S. Multiple stable states in microbial communities explained by the stable marriage problem. The ISME Journal 12, 2823–2834 (2018).

9. May, R. M. Thresholds and breakpoints in ecosystems with a multiplicity of stable states. Nature 269, 471–477 (1977).

10. Critical Transitions in Nature and Society - Scheffer, Marten: 9780691122038 - AbeBooks. https://www.abebooks.fr/9780691122038/Critical-Transitions-Nature-Society-Scheffer-0691122032/plp.

11. Connell, J. H. & Sousa, W. P. On the Evidence Needed to Judge Ecological Stability or Persistence. The American Naturalist 121, 789–824 (1983).

12. Estrela, S., Diaz-Colunga, J., Vila, J. C. C., Sanchez-Gorostiaga, A. & Sanchez, A. Diversity begets diversity under microbial niche construction. 2022.02.13.480281 Preprint at 10.1101/2022.02.13.480281 (2022).

13. Ratzke, C., Denk, J. & Gore, J. Ecological suicide in microbes. Nat Ecol Evol 2, 867–872 (2018).

14. Cremin, K., Duxbury, S. J. N., Rosko, J. & Soyer, O. S. Formation and emergent dynamics of spatially organized microbial systems. Interface Focus 13, 20220062 (2023).

15. Gralka, M., Szabo, R., Stocker, R. & Cordero, O. X. Trophic Interactions and the Drivers of Microbial Community Assembly. Current Biology 30, R1176–R1188 (2020).

16. Scott, M. & Hwa, T. Shaping bacterial gene expression by physiological and proteome allocation constraints. Nat Rev Microbiol 21, 327–342 (2023).

17. Balakrishnan, R., de Silva, R. T., Hwa, T. & Cremer, J. Suboptimal resource allocation in changing environments constrains response and growth in bacteria. Molecular Systems Biology 17, e10597 (2021).

18. Evans, C. R., Kempes, C. P., Price-Whelan, A. & Dietrich, L. E. P. Metabolic Heterogeneity and Cross-Feeding in Bacterial Multicellular Systems. Trends in Microbiology 28, 732–743 (2020).

19. Mukherjee, A. et al. Coexisting ecotypes in long-term evolution emerged from interacting trade-offs. Nat Commun 14, 3805 (2023).

20. Wang, X., Xia, K., Yang, X. & Tang, C. Growth strategy of microbes on mixed carbon sources. Nat Commun 10, 1279 (2019).

21. Perrin, E. et al. Diauxie and co-utilization of carbon sources can coexist during bacterial growth in nutritionally complex environments. Nat Commun 11, 3135 (2020).

22. Monod, J. (1910-1976) A. Recherches Sur La Croissance Des Cultures Bactériennes. (Hermann. Paris, 1941).

23. Amarnath, K. et al. Stress-induced metabolic exchanges between complementary bacterial types underly a dynamic mechanism of inter-species stress resistance. Nat Commun 14, 3165 (2023).

24. Liu, B. et al. Starvation responses impact interaction dynamics of human gut bacteria Bacteroides thetaiotaomicron and Roseburia intestinalis. ISME J 17, 1940–1952 (2023).

25. Malik, A. A. et al. Defining trait-based microbial strategies with consequences for soil carbon cycling under climate change. The ISME Journal 14, 1–9 (2020).

26. Duncan, S. H., Louis, P., Thomson, J. M. & Flint, H. J. The role of pH in determining the species composition of the human colonic microbiota. Environmental Microbiology 11, 2112–2122 (2009).

27. van de Velde, C. et al. Technical versus biological variability in a synthetic human gut community. Gut Microbes 15, 2155019 (2023).

28. Lahti, L., Salojärvi, J., Salonen, A., Scheffer, M. & de Vos, W. M. Tipping elements in the human intestinal ecosystem. Nat Commun 5, 4344 (2014).

29. Lopes, W., Amor, D. R. & Gore, J. Cooperative growth in microbial communities is a driver of multistability. Nat Commun 15, 4709 (2024).

30. Ding, T. & Schloss, P. D. Dynamics and associations of microbial community types across the human body. Nature 509, 357–360 (2014).

31. Moraïs, S. & Mizrahi, I. The Road Not Taken: The Rumen Microbiome, Functional Groups, and Community States. Trends in Microbiology 27, 538–549 (2019).

32. Bashan, A. et al. Universality of human microbial dynamics. Nature 534, 259–262 (2016).

33. Vila, J. C. C., Liu, Y.-Y. & Sanchez, A. Dissimilarity–Overlap analysis of replicate enrichment communities. The ISME Journal 14, 2505–2513 (2020).

34. Ansari, A. F., Reddy, Y. B. S., Raut, J. & Dixit, N. M. An efficient and scalable top-down method for predicting structures of microbial communities. Nat Comput Sci 1, 619–628 (2021).

35. Li, C. et al. BEEM-Static: Accurate inference of ecological interactions from cross- sectional microbiome data. PLOS Computational Biology 17, e1009343 (2021).

36. Bucci, V. et al. MDSINE: Microbial Dynamical Systems INference Engine for microbiome time-series analyses. Genome Biology 17, 121 (2016).

37. Rios Garza, D., Gonze, D., Zafeiropoulos, H., Liu, B. & Faust, K. Metabolic models of human gut microbiota: Advances and challenges. Cell Systems 14, 109–121 (2023).

38. Liu, B., Garza, D. R., Saha, P., Zhou, X. & Faust, K. Exploiting gut microbial traits and trade-offs in microbiome-based therapeutics. Nat Rev Bioeng 1–3 (2024).

39. Nguyen, L. K. & Kulasiri, D. On the functional diversity of dynamical behaviour in genetic and metabolic feedback systems. BMC Systems Biology 3, 51 (2009).

40. Monod, J. The Growth of Bacterial Cultures. Annual Review of Microbiology 3, 371–394 (1949).

41. Fischbach, M. A. & Sonnenburg, J. L. Eating For Two: How Metabolism Establishes Interspecies Interactions in the Gut. Cell Host & Microbe 10, 336–347 (2011).

42. Li, H. New strategies to improve minimap2 alignment accuracy. Bioinformatics 37, 4572– 4574 (2021).

43. O’Leary, N. A. et al. Reference sequence (RefSeq) database at NCBI: current status, taxonomic expansion, and functional annotation. Nucleic Acids Research 44, D733–D745 (2016).

44. Stoddard, S. F., Smith, B. J., Hein, R., Roller, B. R. K. & Schmidt, T. M. rrnDB: improved tools for interpreting rRNA gene abundance in bacteria and archaea and a new foundation for future development. Nucleic Acids Research 43, D593–D598 (2015).

45. Hernandez-Sanabria, E. et al. In vitro Increased Respiratory Activity of Selected Oral Bacteria May Explain Competitive and Collaborative Interactions in the Oral Microbiome. Frontiers in Cellular and Infection Microbiology 7, (2017).

