## Supplementary Text S1 for "Discovery of alternative stable states in a synthetic human gut microbial community"

Here we present the basic components of our kinetic model designed to simulate the flexible life-history strategies of human gut bacteria. The model is based on experiments conducted in monocultures and cocultures of a synthetic community of three species (main Figs. 1 and 2A). Please note that this description is aimed to help readers understand the model’s rationale. A computational implementation of the model and code to reproduce the manuscript figures is available at: https://github.com/danielriosgarza/hungerGamesModel

**Generic model**

*Subpopulations*

Bacteria dynamically change their environments by consuming nutrients and excreting metabolic by-products. Within a community, species are often subjected to changing environments. For example, gut bacteria can be challenged by their host's feeding habits and circadian rhythms, as well as by the metabolic byproducts of neighboring species. To cope with these variations, cells within a population can alter their physiology, leading to phenotypic heterogeneity and the emergence of subpopulations within a single species, which may or may not coexist in space and time.

Subpopulations arise due to changes in cellular physiology in response to environmental variations (either triggered by intrinsic cellular programs or, often, as an unavoidable consequence of being propelled towards intolerable conditions). In our model, we represent these subpopulations as distinct kinetic functions for a given species, each having its own set of kinetic parameters. These functions are linked through transition rates and transition functions, which encode the likelihood that cells transition from one subpopulation to another in response to environmental cues.

*Transition rates (*$r$*)*

Cellular transitions can result from a single process, multiple independent processes, or multiple dependent processes. These transitions may occur rapidly, affecting cells concurrently, or they may take place gradually, leading to the formation of co-existing subpopulations. The rate at which a population transitions from one state to another is modulated by the transition rate ($r$).

*Transition functions (*$Z$*)*

In our model, transition functions encode the environmental cues that trigger transitions between subpopulations. These functions operate like sensors, responding to changes in factors such as pH, metabolite concentrations, or other environmental variables. Transitions can occur independently or involve activation and inhibition mechanisms.

*Independent transitions*

These transitions occur at a constant rate, unaffected by any environmental conditions. In this case, $Z$ equals one, indicating a fixed transition rate. This transition is commonly assumed in scenarios where there is a constant death rate.

The following equation exemplifies an independent transition rate. Here, subpopulation $A$ contributes towards changes in subpopulation $B$ over time, given the counts $N_{A}$ and $N_{B}$ for each subpopulation, respectively:

$$\frac{dN_{B}}{dt}=\left( \underset{N_{A}\to N_{B}}{r} \right)N_{A}+\left( ... \right)$$

The first term on the right-hand side accounts for the contribution of subpopulation $A$ to the growth of subpopulation $B$, where $r$ is the transition rate from subpopulation $A$ to subpopulation $B$. The ellipsis $\left( ... \right)$ represents any additional terms that may affect the time derivative of subpopulation $B$, such as its growth rates or the reactor's dilution rate.

*Activation-like transition*

An activation-like transition is used when a specific molecule or condition triggers the transition between different subpopulations. This transition occurs when the concentration of the molecule or the strength of the condition increases to a certain threshold. For example, a subpopulation of cells in a slow growth mode (subpopulation A) may transition to a fast growth mode (subpopulation B) when the concentration of a metabolite, such as glucose, increases in the environment, and the subpopulation can sense it in a concentration-dependent manner (measured by $S_{M}$).

We use a Hill function^37^ to modulate the transition rate based on the metabolite concentration. The Hill function is a sigmoidal curve that varies from 0 to 1 and represents the fraction of cells that transition from subpopulation A to subpopulation B at a given metabolite concentration. The transition rate is multiplied by the Hill function to obtain the effective transition rate, which is then used to compute the contribution towards the derivative of subpopulation B.

$$\underset{N_{A}\to N_{B}}{Z}=\frac{S_{M}^{h}}{K^{h}+S_{M}^{h}}$$

Here, $K$ is the half-max constant, which is in the same unit as $S_{M}$ (e.g., $mM$ if we are talking about a metabolite). It defines the concentration where the transition function equals to $0.5$ (the subpopulation $A$ transitions to $B$ at half the transition rate).

The parameter $h$ is the Hill coefficient that determines how steep the transition will occur around the half-max constant (i.e., its slope).

The contribution to the derivative is then:

$$\frac{dN_{B}}{dt}=\left( \frac{S_{M}^{h}}{K^{h}+S_{M}^{h}} \right)\left( \underset{N_{A}\to N_{B}}{r} \right)N_{A}+\left( ... \right)$$

The model can also support a pure logical rule instead of a Hill function (which is equivalent to a Hill function with an infinite exponent), for instance:

$$\begin{matrix} & \underset{\left( N_{A}\to N_{B} \right)}{Z}=\left\{ \begin{matrix} 1, & \text{if }S_{M}>\text{ some value} \\ & \\ 0, & \text{otherwise} \end{matrix} \right. & \end{matrix}$$

This can be used initially to fit the model to data (explained below), as it requires less parameters. It can then easily be replaced by the smooth activation function described above.

*Inhibition*

Inhibition occurs when the transition is triggered by the 'lack' or 'depletion' of something. For example, when the depletion of a metabolite $M$ that is used as an energy source leads to an increase in cell death rate, which is represented by an increase in the rate at which cells in subpopulation A (live) transition to subpopulation B (dead). The transition function used for inhibition is similar to the activation function, but is inverter:

$$\underset{N_{A}\to N_{B}}{Z}=\frac{K^{h}}{K^{h}+S_{M}^{h}}$$

The pure logical rule would be:

$$\begin{matrix} & \underset{\left( N_{A}\to N_{B} \right)}{Z}=\left\{ \begin{matrix} \underset{N_{A}\to N_{B}}{r}, & \text{if }S_{M}<\text{ some value} \\ & \\ 0, & \text{otherwise} \end{matrix} \right. & \end{matrix}$$

Note that other logical rules (e.g., if, if-else statements) as functions of the environment (pH or metabolite concentrations) are also supported by our model.

*Metabolites*

Species interactions and survival within an environment are largely shaped by the production and consumption of metabolites. In our model, we identified some of the nutrients available in the growth medium we used, namely the Wilkins-Chalgren anaerobic broth. This medium contains glucose and pyruvate as added carbon sources. In addition, we measured the time-series concentration of trehalose, a disaccharide produced by yeast and introduced into the medium through yeast extract. We also confirmed the presence of glutamate and mannose through single measurements (not time series).

Based on these observations and certain justified assumptions, we included in the model the concentration of several compounds, including glucose, pyruvate, trehalose, lactate, acetate, succinate, butyrate, formate, glutamate, and mannose. These compounds are consumed or produced by at least one of the three species in our model, as we explained in the model state equations (described below).

*Feeding terms*

Active subpopulations utilize nutrients for growth and maintenance, leading to the secretion of by-products. This process can be conceptualized as a directed graph, with nodes representing metabolites and edges representing biochemical reactions that transform one metabolite into another. The graph initiates with nutrients present in the medium and follows the flow of metabolites through the network until the by-products are secreted back into the medium, as illustrated below:

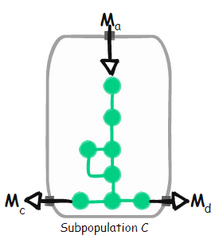

Mathematically, the consumption of metabolite $M_{a}$ and production of metabolites $M_{d}$ and $M_{c}$ are connected by the consumption rate of metabolite $M_{a}$. For consumption, we employ the Monod equation^38^. From here on we denote to the concentration of metabolites measured in $mM$ and the subpopulation counts measured in ${10}^{-5}cells\mu L^{-1}$ by their identifying names.

$$\frac{dM_{a}}{dt}=-\gamma_{C,M_{a}}\left( \frac{M_{a}}{K_{\left( C,M_{a} \right)}+M_{a}} \right)C\mu max_{C}+\left( ... \right)$$

$$\frac{dM_{d}}{dt}=\gamma_{C,M_{d}}\left( \frac{M_{a}}{K_{\left( C,M_{a} \right)}+M_{a}} \right)C\mu max_{C}+\left( ... \right)$$

$$\frac{dM_{c}}{dt}=\gamma_{C,M_{c}}\left( \frac{M_{a}}{K_{\left( C,M_{a} \right)}+M_{a}} \right)C\mu max_{C}+\left( ... \right)$$

Where $\gamma_{\text{subpopulation},\text{metabolite}}$ represent the relative contribution (weight) of the subpopulation growth towards the consumption (if negative) or production (if positive) of the respective metabolite, and $K$ is the Monod constant. $\mu max_{C}$ is the maximum growth rate reached by the subpopulation $x_{C}.$

In our model, we represented each species' core energy metabolism as an independent subnetwork extracted from its respective genome-scale metabolic model. We focused on the fraction of the network that could explain the changes in metabolite concentrations measured during growth in monocultures. This approach was supported by the analysis of gene expression and differential expression of metabolic genes (see Fig. 1 in the main text).

Analyzing the core energy graphs allows us to link subpopulations to sets of metabolites that contribute to their growth. These sets of metabolites are represented as feeding terms, which can display either an "or" (additive) relationship or an "and" (multiplicative) relationship between metabolites, based on the topology of the metabolic network.

Analyzing the core energy graphs allows us to link subpopulations to sets of metabolites that contribute to their growth. These sets of metabolites are represented as feeding terms, which can display either an "or" (additive) relationship or an "and" (multiplicative) relationship between metabolites, based on the topology of the metabolic network.

*AND*

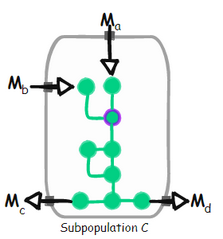

The network requires the influx of both $M_{a}$ and $M_{b}$ to function (otherwise the intracellular metabolite that is highlighted in purple cannot be produced)

$$\frac{dM_{a}}{dt}=-\gamma_{C,M_{a}}\left( \frac{M_{a}}{K_{C,M_{a}}+M_{a}} \right)\left( \frac{M_{b}}{K_{C,M_{b}}+M_{b}} \right)C\mu max_{C}+\left( ... \right)$$

$$\frac{dM_{b}}{dt}=-\gamma_{C,M_{b}}\left( \frac{M_{a}}{K_{C,M_{a}}+M_{a}} \right)\left( \frac{M_{b}}{K_{C,M_{b}}+M_{b}} \right)C\mu max_{C}+\left( ... \right)$$

$$\frac{dM_{c}}{dt}=\gamma_{C,M_{c}}\left( \frac{M_{a}}{K_{C,M_{a}}+M_{a}} \right)\left( \frac{M_{b}}{K_{C,M_{b}}+M_{b}} \right)C\mu max_{C}+\left( ... \right)$$

$$\frac{dM_{d}}{dt}=\gamma_{C,M_{d}}\left( \frac{M_{a}}{K_{C,M_{a}}+M_{a}} \right)\left( \frac{M_{b}}{K_{C,M_{b}}+M_{b}} \right)C\mu max_{C}+\left( ... \right)$$

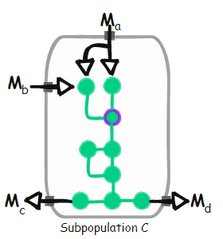
*Boost*

$M_{b}$ may increase the flux through the network, but only in the presence of $M_{a}$. If $M_{b}$ is absent, the network can still be functional, provided thar $M_{a}$ is present, but not the other way around.

$$\frac{dM_{a}}{dt}=-\gamma_{C,M_{a}}\left[ \left( \frac{M_{a}}{K_{C,M_{a}}+M_{a}} \right)+\left( \frac{M_{a}}{K_{C,M_{a}}+M_{a}} \right)\left( \frac{M_{b}}{K_{C,M_{b}}+M_{b}} \right) \right]C\mu max_{C}+\left( ... \right)$$

$$\frac{dM_{b}}{dt}=-\gamma_{C,M_{b}}\left[ \left( \frac{M_{a}}{K_{C,M_{a}}+M_{a}} \right)+\left( \frac{M_{a}}{K_{C,M_{a}}+M_{a}} \right)\left( \frac{M_{b}}{K_{C,M_{b}}+M_{b}} \right) \right]C\mu max_{C}+\left( ... \right)$$

$$\frac{dM_{c}}{dt}=\gamma_{C,M_{c}}\left[ \left( \frac{M_{a}}{K_{\left( M_{a},C \right)}+M_{a}} \right)+\left( \frac{M_{a}}{K_{\left( M_{a},C \right)}+M_{a}} \right)\left( \frac{M_{b}}{K_{\left( M_{b},C \right)}+M_{b}} \right) \right]C\mu max_{C}+\left( ... \right)$$

$$\frac{dM_{d}}{dt}=\gamma_{C,M_{d}}\left[ \left( \frac{M_{a}}{K_{C,M_{a}}+M_{a}} \right)+\left( \frac{M_{a}}{K_{C,M_{a}}+M_{a}} \right)\left( \frac{M_{b}}{K_{C,M_{b}}+M_{b}} \right) \right]C\mu max_{C}+\left( ... \right)$$

*Environment (reactor)*

Environmental conditions substantially influence microbial growth and interactions. Changes in the environment can lead to shifts in both the taxonomic composition and the function of the microbial system. A common approach to understanding these impacts is through collecting experimental data under varying conditions. However, we can take this a step further by encoding our knowledge into a dynamic model that explicitly accounts for environmental conditions. Once calibrated with parameters that best fit the experimental data, this model helps us predict the impact of environmental conditions on our system. These predictions can be validated through further experiments, and if the results diverge, they serve to highlight key knowledge gaps.

Our model considers the impact of different environmental conditions by encoding the system's nutrient flows and pH dynamics. Specifically, we consider the consumption and production of metabolites by growing subpopulations combined with their responses to environmental changes. The composition of the environment affects the pH, which, in turn, impacts the growth and survival of the different subpopulations.

*Environment pH*

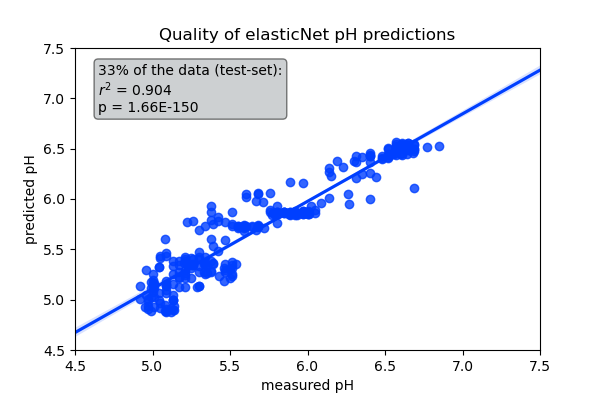
To predict pH changes in the environment, we developed an elastic net model using paired measurements of pH and fermentation acid concentrations. The model considers the concentrations of lactate, acetate, formate, and butyrate to estimate the environmental pH. We trained it using a randomly selected subset constituting 67% of all our experiments, encompassing both monocultures and cocultures. We evaluated the model's performance by comparing predicted pH values with actual measurements from the remaining 33% of the test cases, which were not part of the model training process.
*pH limitation*

In our model, the pH value at each time point is determined by the elastic net model, which is based on the concentrations of fermentation acids. The pH varies as the subpopulations consume and produce these acids. The fluctuating pH influences the growth rates of the subpopulations, according to their preferred pH and their sensitivity to pH variations. It is important to note that our model is only concerned with pH fluctuations below the initial anaerobic pH of the system (approximately 6.7), as we did not observe any pH increases beyond this value.

To model the impact of pH on growth, we used the probability density function of the gamma distribution, with some modifications. The gamma distribution is commonly used in statistics to model positive random variables with a skewed distribution. Since we aim to multiply the pH sensitivity by the growth rate, we require a function (referred to as $\xi_{A}\left( pH \right)$ ) that takes the current pH as input and returns a value between zero and one. This output depends on the subpopulation's sensitivity to deviations from its preferred pH. The function must output a value of one at the optimal pH (or zero if we refer to pH sensitivity, where sensitivity = $1-\xi_{A}\left( pH \right)$ ).

Starting from the probability distribution function of the gamma distribution:

$$\varphi\left( pH;\alpha,\beta\right)=\frac{\beta^{\alpha}}{\Gamma\left( \alpha\right)}\left[ pH \right]^{\alpha-1} e^{-\beta\left[ ph \right]}$$

where $\alpha$ and $\beta$ are the shape and rate parameters of the gamma distribution, and $\Gamma$ is the gamma function. To obtain the properties we desire, we reparameterized the function using the optimal pH instead of the rate (𝛽), and we adjusted the function to have a maximum value of exactly one. To achieve this, we set the mode (𝑚) of the gamma distribution to be the optimal pH and solved for $\beta$:

$$m=\frac{\alpha-1}{\beta}$$

$$pHopt=\frac{\alpha-1}{\beta}$$

$$\beta=\frac{\alpha-1}{pHopt}$$

Next, to ensure that the optimal pH returns a value of one, we divide the function by its maximum value. Note that the maximum value of the function occurs exactly at the optimal pH. Tying it together, the impact of pH on the growth of subpopulation A is modeled by:

$$\xi_{A}\left( pH;pHopt_{A},\alpha_{A} \right)=\frac{1}{\varphi\left( pHopt_{A};\alpha_{A},\left( \frac{\alpha_{A}-1}{pHopt_{A}} \right) \right)}\varphi\left( pH;\alpha_{A},\left( \frac{\alpha_{A}-1}{pHopt_{A}} \right) \right)$$

($\varphi$ is defined above)

Below, we show the pH sensitivity of a subpopulation that has $pHopt=7.0$ for different values of $\alpha$ (note that $\alpha$ needs to be greater than one):

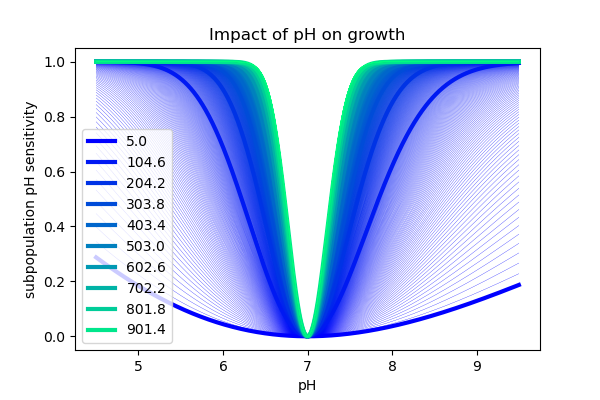

*Growth*

We now have all the ingredients for our general growth model. Our general growth model comprises a kinetic equation for each subpopulation, describing changes in its concentration in units of ${10}^{-5}cells\mu L^{-1}$, comparable to flow cytometer counts. These growth equations have four components:

**a) Maximum intrinsic growth rate (**$\boldsymbol{\mu max}$**)**

This represents the maximum growth rate that a subpopulation's physiology supports. In the absence of any limiting factors, cells would grow at a rate equal to $\mu max$.

**b) pH limitation (**$\boldsymbol{\xi}$**)**

This component considers the impact of pH on growth, using a modified probability density function of the gamma distribution. The function $\xi_{A}\left( pH \right)$ returns a value between zero and one, depending on how sensitive the subpopulation is to deviations from its preferred pH.

**c) Nutrient consumption**

This is based on the additive combination of the feeding terms that are actively used by a subpopulation.

**d) Subpopulation transitions**

This component considers incoming and outgoing subpopulation transitions.

Next, we illustrate the model equations for the following toy diagram where subpopulation $A$ grows on $M_{1}$ and secretes $M_{2}$. When $M_{1}$ is depleted from the media, if both $M_{3}$ and $M_{4}$ are available, it switches to a new growth mode, summarized by subpopulation $B$ that grows by simultaneously consuming $M_{3}$ and $M_{4}$ while still secreting $M_{2}$.

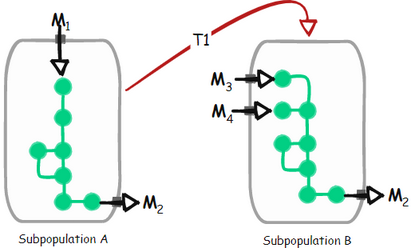

*Growth of subpopulation A*

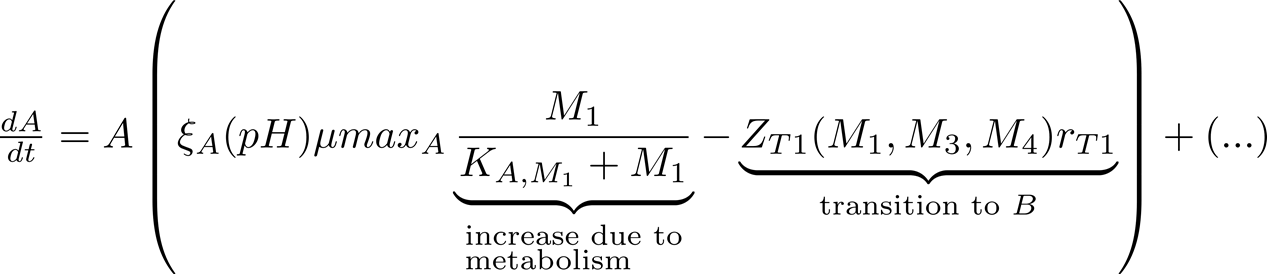

$\xi_{A}\left( pH \right)$ is explained in the “pH limitation” session above.

The transition function is:

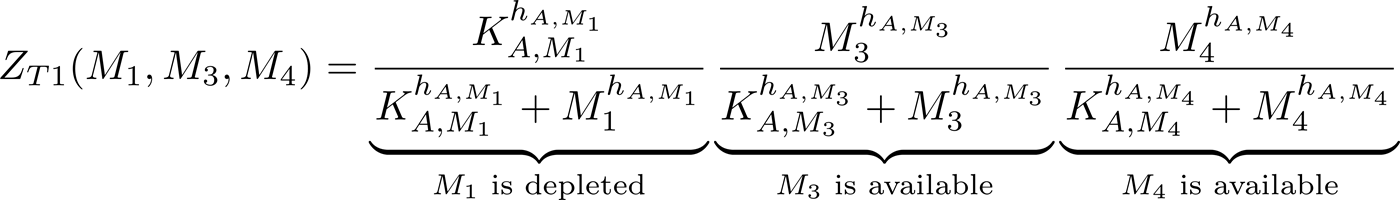

Growth of subpopulation $B$

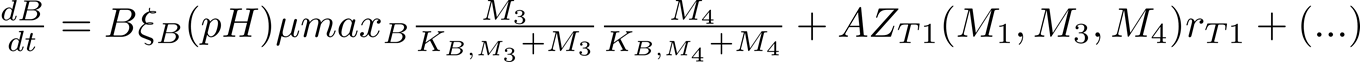

Consumption of metabolite $M_{1}$

$$\frac{dM_{1}}{dt}=-\gamma_{A,M_{1}}\left( \frac{M_{1}}{K_{A,M_{1}}+M_{1}} \right)A\mu max_{A}+\left( ... \right)$$

Consumption of metabolite $M_{3}$

$$\frac{dM_{3}}{dt}=-\gamma_{B,M_{3}}\left( \frac{M_{3}}{K_{B,M_{3}}+M_{3}} \right)\left( \frac{M_{4}}{K_{B,M_{4}}+M_{4}} \right)B\mu max_{B}+\left( ... \right)$$

Consumption of metabolite $M_{4}$

$$\frac{dM_{4}}{dt}=-\gamma_{B,M_{4}}\left( \frac{M_{3}}{K_{B,M_{3}}+M_{3}} \right)\left( \frac{M_{4}}{K_{B,M_{4}}+M_{4}} \right)B\mu max_{B}+\left( ... \right)$$

Production of metabolite $M_{2}$

$$\frac{dM_{2}}{dt}=\gamma_{A,M_{2}}\left( \frac{M_{1}}{K_{A,M_{1}}+M_{1}} \right)A\mu max_{A}+\gamma_{B,M_{2}}\left( \frac{M_{3}}{K_{B,M_{3}}+M_{3}} \right)\left( \frac{M_{4}}{K_{B,M_{4}}+M_{4}} \right)B\mu max_{B}+\left( ... \right)$$

pH

$$pH=\beta M_{2}$$

where $\beta$ is the elastic net weight attributed to $M_{2}$.

*Pulses*

In addition to subpopulations growing by the exchange of metabolites and being influenced by environmental pH, the community is also affected by the environmental regime, particularly by the way matter flows in and out of the system. In our model, the environmental regime is controlled by subdividing the simulation into arbitrary time intervals, which we refer to as "pulses." Within a pulse, the following events are supported:

1. **Noncontinuous inflow (**$\boldsymbol{v}_{\boldsymbol{in}}$**) or outflow (**$\boldsymbol{v}_{\boldsymbol{out}}$**) of volume:**

This is performed in a single step at the beginning of the pulse. Multiple pulses are required to simulate multiple influx and outflux events, as in a serial passage experiment. The content of the influx volume, defined by the user, could include fresh metabolites or even spent media from another culture. The user may also set its pH and, if necessary, a specific concentration of bacteria to simulate migration.

2. **Continuous inflow (**$\boldsymbol{q}_{\boldsymbol{in}}$**) and outflow (**$\boldsymbol{q}_{\boldsymbol{out}}$**) of volume per time unit**:

Volume is added to or removed from the reactor continuously during a pulse. Similar to the non-continuous events, the user defines the metabolome, microbiome, and pH of the feed.

With these events, a wide range of community regimes can be simulated. For example, one may start a pulse in a batch culture, then transition to a chemostat with a nutrient-rich medium, and then move back to a batch culture. Simulations could include serial passages or even mass transfer between separate reactors, allowing the exploration of regimes that lead to multistability and trigger alternative community states.

*State equations*

Here, we define the state variables and equations that simulate bacterial growth and life strategies in our three-species synthetic community (depicted in Figure 2A of the main text). The model components are assigned according to the description of the generic model explained above and informed by our investigation of the growth kinetics of each species (see Figure 1 in the main text). We assigned parameters that represent the best fit to three independent monoculture experiments, each with two or more biological replicates.

Some of the notable features that we found while investigating the physiology of our bacteria in WC media in experiments were incorporated into the model and are summarized below (also see Figure 1 of the main text):

*Blautia hydrogenotrophica* DSM 10507 (referred to as "Bh").

- Bh shows preference for trehalose over glucose

When both trehalose ($s_{1}$) and glucose ($s_{3}$) are present in the medium, Bh first consumes trehalose and neglects glucose until trehalose is depleted.

Bh's genome contains a trehalose-specific PTS gene, which is found to be overexpressed when Bh grows on trehalose. Interestingly, the genome does not contain the glucose-specific IIA component of the PTS system gene that is found in closely-related *Blautia* and *Ruminococcus* strains. In the presence of trehalose, the non-PTS glucose transporter is inhibited while the trehalose-specific PTS gene is actively expressed. To capture this behavior in our model, we used the transition function ($Z_{1}$), which triggers a switch from a subpopulation ($x_{a}$) that does not uptake glucose to a subpopulation that uptakes glucose ($x_{b}$) based on the concentration of trehalose.

We found that adding a higher concentration of trehalose to the media ($4mM$) completely prevented Bh from shifting to glucose consumption. This behavior was not observed when increasing the concentration of pyruvate.

We also found that increasing trehalose leads to an increase in lactate production, while increasing pyruvate results in an increase in acetate, but not lactate production.

In standard WC medium, pyruvate was depleted before glucose was consumed. However, when we supplemented the medium with pyruvate, we observed co-consumption of glucose and pyruvate. In contrast, the presence of trehalose inhibits the uptake of glucose.

- Bh exhibits higher growth rates on glucose compared to trehalose and pyruvate

We observed higher growth-rates during the glucose-consuming stage compared to the trehalose-consuming stage.

- Bh exhibits glucose co-limitation

Not all of the glucose is consumed from the media before the culture enters the stationary and death phases, suggesting co-limitation with another substrate. The core-metabolic pathway suggests that growth on glucose would be favored by glutamate fermentation, which would provide an additional mol of CO2 and a reduced ferredoxin that could be used to pump protons through the RNF system. This favors ATP production through the membrane ATPase (ATPS4), which is driven by a proton gradient. We modeled this behavior as a co-consumption of glucose and glutamate as the genes for glutamate fermentation are significantly overexpressed during growth on glucose. We confirmed glutamate depletion from the spent media by measuring the levels of amino acids before and after fermentation.

*Bacteroides thetaiotaomicron* VPI-5482 (referred to as "Bt")

- Bt produces a range of fermentation acids that significantly decrease the medium pH

In our experiments, Bt was observed to rapidly consume glucose and pyruvate, while producing a variety of fermentation acids. This resulted in a swift drop in the medium's pH.

- Bt is inhibited by low pH values

Bt is known to be sensitive to low pH levels^26^, a fact we corroborated by incubating the cells across different pH ranges. These experiments revealed that while the population did not grow at pH levels < 5, most cells remained viable. As such, we modelled pH's impact as a growth inhibitor.

But we also found that when carbon sources are exhausted most Bt cells lose their viability. This was confirmed through assessing cell permeability with PI staining. To represent this in our model, we introduced transition functions from active to inactive subpopulations that are triggered by nutrient depletion at low pH.

- Bt fixes CO2, producing succinate

Bt fixes CO2 by converting phosphoenolpyruvate (a C3 molecule) into oxaloacetate (a C4 molecule) in a process that mirrors carbon fixation in plants. Via the reductive carboxylation of phosphoenolpyruvate^39^, Bt generates ATP and produces succinate.

- Bt shows a second growth peak in WC medium

We consistently observed a second growth peak before the majority of the cell population transitioned to an inactive state. We attributed this second peak to the consumption of mannose, which is present in low concentrations in the complex medium. Mannose depletion was confirmed through single-point measurements and suggested by the gene expression data.

*Roseburia intestinalis* L1-82 (referred to as "Ri")

- Ri is a butyrate producer

By studying its core metabolic pathway, we found that Ri produces butyrate through the reverse beta oxidation pathway. In our experiments, Ri quickly consumed glucose and pyruvate and produced butyrate, acetate, and lactate.

- Ri has lesser impact on the medium pH

Unlike Bt, Ri exerts a weaker effect on the pH of the medium despite its high growth rate.

- Ri enters a slow growth mode characterized by the consumption of lactate and acetate

We observed that some of the lactate and acetate that are produced during growth in glucose and pyruvate, later get consumed, leading to a gradual increase in butyrate.

Following the consumption of glucose and pyruvate and production of butyrate, lactate, and acetate, most cells burst and are no longer detected by flow cytometry. However, a subset of cells can persist for several days, possibly entering a slow growth mode^24^. We modeled this behavior by having cells quickly die in the absence of glucose and transition to a slow growth mode when triggered by lactate.

| **States**  Symbol | type | state | units |
| --- | --- | --- | --- |
| $x_{a}$ | subpopulation | Bh subpopulation that consumes trehalose | $\frac{{10}^{5}\text{cells}}{\mu L}$ |
| $x_{b}$ | subpopulation | Bh subpopulation that consumes glucose | $\frac{{10}^{5}\text{cells}}{\mu L}$ |
| $x_{c}$ | subpopulation | Bh inactive subpopulation | $\frac{{10}^{5}\text{cells}}{\mu L}$ |
| $x_{d}$ | subpopulation | Bh dead subpopulation | $\frac{{10}^{5}\text{cells}}{\mu L}$ |
| $x_{e}$ | subpopulation | Bt subpopulation that consumes glucose | $\frac{{10}^{5}\text{cells}}{\mu L}$ |
| $x_{f}$ | subpopulation | Bt subpopulation that consumes mannose | $\frac{{10}^{5}\text{cells}}{\mu L}$ |
| $x_{g}$ | subpopulation | Bt inactive subpopulation | $\frac{{10}^{5}\text{cells}}{\mu L}$ |
| $x_{h}$ | subpopulation | Bt dead subpopulation | $\frac{{10}^{5}\text{cells}}{\mu L}$ |
| $x_{i}$ | subpopulation | Ri subpopulation that consumes glucose | $\frac{{10}^{5}\text{cells}}{\mu L}$ |
| $x_{j}$ | subpopulation | Ri subpopulation that consumes lactate and acetate | $\frac{{10}^{5}\text{cells}}{\mu L}$ |
| $x_{k}$ | subpopulation | Ri inactive subpopulation | $\frac{{10}^{5}\text{cells}}{\mu L}$ |
| $x_{l}$ | subpopulation | Ri dead subpopulation | $\frac{{10}^{5}\text{cells}}{\mu L}$ |
| $s_{1}$ | metabolite | trehalose | $mM$ |
| $s_{2}$ | metabolite | pyruvate | $mM$ |
| $s_{3}$ | metabolite | glucose | $mM$ |
| $s_{4}$ | metabolite | glutamate | $mM$ |
| $s_{5}$ | metabolite | lactate | $mM$ |
| $s_{6}$ | metabolite | acetate | $mM$ |
| $s_{7}$ | metabolite | mannose | $mM$ |
| $s_{8}$ | metabolite | succinate | $mM$ |
| $s_{9}$ | metabolite | formate | $mM$ |
| $s_{10}$ | metabolite | butyrate | $mM$ |
| pH | pH | potential of hydrogen |  |

Equations

$$\boldsymbol{x}_{\boldsymbol{a}}$$

| diagram | state | unit | description |
| --- | --- | --- | --- |
| 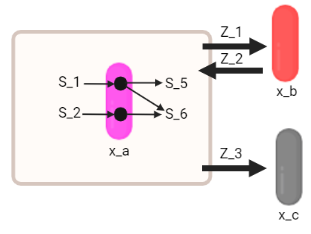 | $x_{a}$ | $\frac{{10}^{5}\text{cells}}{\mu L}$ | live Bh subpopulation that grows on trehalose ($s_{1}$) or pyruvate ($s_{2}$), and produces acetate ($s_{6}$) and lactate ($s_{5}$) |

$$\frac{dx_{a}}{dt}=x_{a}\left[ \xi_{x_{a}}\mu max_{x_{a}}\left( \frac{s_{1}}{s_{1}+K_{x_{a},s_{1}}}+\frac{s_{2}}{s_{2}+K_{x_{a},s_{2}}} \right)-\left( Z_{1}+Z_{3} \right) \right]+x_{b}Z_{2}$$

$$\xi_{x_{a}}\left( pH \right)=\frac{1}{\varphi\left( p_{x_{a}};\alpha_{x_{a}},\frac{\alpha_{x_{a}}-1}{p_{x_{a}}} \right)}\varphi\left( pH;\alpha_{x_{a}},\frac{\alpha_{x_{a}}-1}{p_{x_{a}}} \right)$$

The $\varphi$ function is explained in the “pH limitation” section of the generic model.

transition from $x_{a}$ to $x_{b}$, triggered by low trehalose concentration ($s_{1}$):

$$Z_{1}=\left( \frac{l_{1,s_{1}}^{h_{1,s_{1}}}}{S_{1}^{h_{1,s_{1}}}+l_{1,s_{1}}^{h_{1,s_{1}}}} \right)r_{1}$$

transition from $x_{b}$ to $x_{a}$ triggered by high trehalose concentration ($s_{1}$):

$$Z_{2}=\left( \frac{S_{1}^{h_{2,s_{1}}}}{S_{1}^{h_{2,s_{1}}}+l_{2,s_{1}}^{h_{2,s_{1}}}} \right)r_{2}$$

transition from $x_{a}$ to $x_{c}$:

$$Z_{3}=r_{3}$$

$$\boldsymbol{x}_{\boldsymbol{b}}$$

| diagram | state | unit | description |
| --- | --- | --- | --- |
| 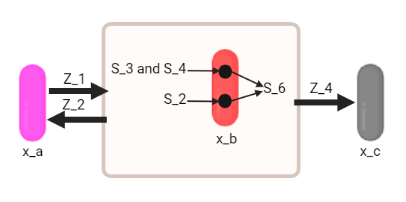 | $x_{b}$ | $\frac{{10}^{5}\text{cells}}{\mu L}$ | live Bh subpopulation that grows on glucose ($s_{3}$) and glutamate ($s_{4}$), and produces acetate ($s_{6}$) |

$$\frac{dx_{b}}{dt}=x_{b}\left[ \xi_{x_{b}}\mu max_{x_{b}}\left( \frac{s_{3}}{s_{3}+K_{x_{b},s_{3}}}\frac{s_{4}}{s_{4}+K_{x_{b},s_{4}}}+\frac{s_{2}}{s_{2}+K_{x_{b},s_{2}}} \right)-\left( Z2+Z_{4} \right) \right]+x_{a}Z_{1}$$

$$\xi_{x_{b}}\left( pH \right)=\frac{1}{\varphi\left( p_{x_{b}};\alpha_{x_{b}},\frac{\alpha_{x_{b}}-1}{p_{x_{b}}} \right)}\varphi\left( pH;\alpha_{x_{b}},\frac{\alpha_{x_{b}}-1}{p_{x_{b}}} \right)$$

transition from $x_{b}$ to $x_{c}$ triggered by low concentrations of glucose ($s_{3}$) or glutamate ($s_{4}$):

$$Z_{4}=\frac{l_{4,s_{3},s_{4}}^{h_{4,s_{3},s_{4}}}}{\left( \frac{S_{3}+S_{4}\sqrt{\left( S_{3}-S_{4} \right)^{2}}}{2} \right)^{h_{4,s_{3},s_{4}}}+l_{4,s_{3},s_{4}}^{h_{4,s_{3},s_{4}}}}r_{4}$$

$$\boldsymbol{x}_{\boldsymbol{c}}$$

| diagram | state | unit | description |
| --- | --- | --- | --- |
| 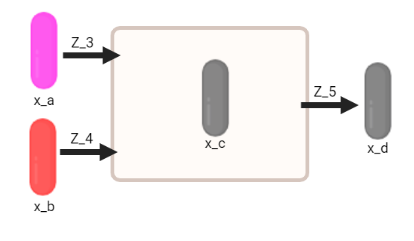 | $x_{c}$ | $\frac{{10}^{5}\text{cells}}{\mu L}$ | Inactive Bh subpopulation that is PI-positive and eventually bursts |

$$\frac{dx_{c}}{dt}=x_{a}Z_{3}+x_{b}Z_{4}-x_{c}Z_{5}$$

$$\frac{dx_{d}}{dt}=x_{c}Z_{5}$$

fixed burst rate:

$$Z_{5}=r_{5}$$

$$\boldsymbol{x}_{\boldsymbol{e}}$$

| diagram | state | unit | description |
| --- | --- | --- | --- |
| 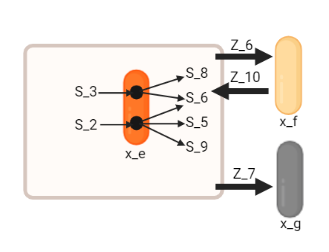 | $x_{e}$ | $\frac{{10}^{5}\text{cells}}{\mu L}$ | live Bt subpopulation that grows on glucose ($s_{3}$) or pyruvate ($s_{2}$), and produces acetate ($s_{6}$), lactate ($s_{5}$), formate ($s_{9}$), and succinate ($s_{8}$) |

$$\frac{dx_{e}}{dt}=x_{e}\left[ \xi_{x_{e}}\mu max_{x_{e}}\left( \frac{s_{2}}{s_{2}+K_{x_{e},s_{2}}}+\frac{s_{3}}{s_{3}+K_{x_{e},s_{3}}} \right)-\left( Z_{6}-Z_{7} \right) \right]+x_{f}Z_{10}$$

$$\xi_{x_{e}}\left( pH \right)=\frac{1}{\varphi\left( p_{x_{e}};\alpha_{x_{e}},\frac{\alpha_{x_{e}}-1}{p_{x_{e}}} \right)}\varphi\left( pH;\alpha_{x_{e}},\frac{\alpha_{x_{e}}-1}{p_{x_{e}}} \right)$$

transition from $x_{e}$ to $x_{f}$ triggered by low glucose ($s_{3}$) and high mannose ($s_{7}$) concentrations:

$$Z_{6}=\left( \frac{l_{6,s_{3}}^{h_{6,s_{3}}}}{s_{3}^{h_{6,s_{3}}}+l_{6,s_{3}}^{h_{6,s_{3}}}} \right)\left( \frac{s_{7}^{h_{6,s7}}}{s_{7}^{h_{6,s7}}+l_{6,s7}^{h_{6,s7}}} \right)r_{6}$$

transition from $x_{e}$ to $x_{g}$ triggered by low glucose concentration and low pH:

$$Z_{7}=\left( \frac{l_{7,s3}^{h_{7,s3}}}{s_{3}^{h_{7,s3}}+l_{7,s3}^{h_{7,s3}}}\frac{l_{7,\text{pH}}^{h_{7,\text{pH}}}}{l_{7,\text{pH}}^{h_{7,\text{pH}}}+\text{pH}^{h_{7,\text{pH}}}} \right)r_{7}$$

transition from $x_{f}$ to $x_{e}$ triggered by high glucose:

$$Z_{10}=\frac{s_{3}^{h_{10},s_{3}}}{s_{3}^{h_{10},s_{3}}+l_{10,s_{3}}^{h_{10,s_{3}}}}$$

$$\boldsymbol{x}_{\boldsymbol{f}}$$

| Diagram | state | unit | description |
| --- | --- | --- | --- |
| 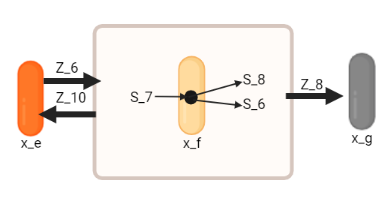 | $x_{f}$ | $\frac{{10}^{5}\text{cells}}{\mu L}$ | live Bt subpopulation that grows on mannose ($s_{7}$), and produces acetate ($s_{6}$) and succinate ($s_{8}$) |

$$\frac{dx_{f}}{dt}=x_{f}\left[ \xi_{x_{f}}\mu max_{x_{f}}\left( \frac{s_{7}}{s_{7}+K_{x_{f},s_{7}}} \right)-\left( Z_{8}+Z_{10} \right) \right]+x_{e}Z_{6}$$

$$\xi_{x_{f}}\left( pH \right)=\frac{1}{\varphi\left( p_{x_{f}};\alpha_{x_{f}},\frac{\alpha_{x_{f}}-1}{p_{x_{f}}} \right)}\varphi\left( pH;\alpha_{x_{f}},\frac{\alpha_{x_{f}}-1}{p_{x_{f}}} \right)$$

transition from $x_{f}$ to $x_{g}$ triggered by low mannose concentration ($s_{7}$) and low pH:

$$Z_{8}=\left( \frac{l_{8,s_{7}}^{h_{8,s_{7}}}}{s_{7}^{h_{8,s_{7}}}+l_{8,s_{7}}^{h_{8,s_{7}}}}\frac{l_{8,\text{pH}}^{h_{8,\text{pH}}}}{l_{8,\text{pH}}^{h_{8,\text{pH}}}+\text{pH}^{h_{8,\text{pH}}}} \right)r_{8}$$

$$\boldsymbol{x}_{\boldsymbol{g}}$$

| diagram | state | unit | description |
| --- | --- | --- | --- |
| 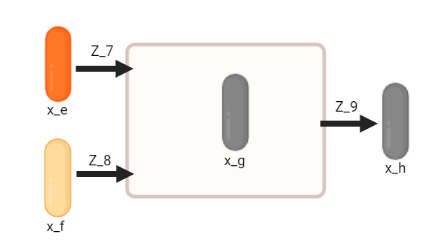 | $x_{g}$ | $\frac{{10}^{5}\text{cells}}{\mu L}$ | inactive Bt subpopulation that is PI-positive and eventually bursts |

$$\frac{dx_{g}}{dt}=x_{e}Z_{7}+x_{f}Z_{8}-x_{g}Z_{9}$$

$$\frac{dx_{h}}{dt}=x_{g}Z_{9}$$

Fixed burst rate:

$Z_{9}=r_{9}$

$$\boldsymbol{x}_{\boldsymbol{i}}$$

| diagram | state | unit | description |
| --- | --- | --- | --- |
| 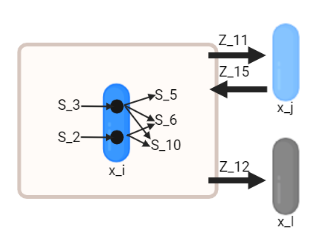 | $x_{i}$ | $\frac{{10}^{5}\text{cells}}{\mu L}$ | live Ri subpopulation that grows on glucose ($s_{3}$) or pyruvate ($s_{2}$), and produces acetate ($s_{6}$), lactate ($s_{5}$), and butyrate ($s_{10}$) |

$$\frac{dx_{i}}{dt}=x_{i}\left[ \xi_{x_{i}}\mu max_{x_{i}}\left( \frac{s_{3}}{s_{3}+K_{x_{i},s_{3}}}+\frac{s_{2}}{s_{2}+K_{x_{i},s_{2}}} \right)-\left( Z_{11}+Z_{12} \right) \right]+x_{j}Z_{15}$$

$$\xi_{x_{i}}\left( pH \right)=\frac{1}{\varphi\left( p_{x_{i}};\alpha_{x_{i}},\frac{\alpha_{x_{i}}-1}{p_{x_{i}}} \right)}\varphi\left( pH;\alpha_{x_{i}},\frac{\alpha_{x_{i}}-1}{p_{x_{i}}} \right)$$

transition from $x_{i}$ to $x_{j}$ triggered by high lactate concentration ($s_{5}$):

$$Z_{11}=\frac{s_{5}^{h_{11,s_{5}}}}{s_{5}^{h_{11,s_{5}}}+l_{11,s_{5}}^{h_{11,s_{5}}}}r_{11}$$

transition from $x_{i}$ to $x_{l}$ triggered by low glucose ($s_{3}$) concentration and pyruvate ($s_{2}$):

$Z_{12}=\frac{l_{12,s_{3},s_{2}}^{h_{12,s_{3},s_{2}}}}{\left( s_{3}+s_{2} \right)^{h_{12,s_{3},s_{2}}}+l_{12,s_{3},s_{2}}^{h_{12,s_{3},s_{2}}}}r_{12}$

transition from $x_{j}$ to $x_{i}$ triggered by high concentration of glucose ($s_{3}$) or pyruvate ($s_{2}$):

$Z_{15}=\frac{\left( s_{3}+s_{2} \right)^{h_{15,s_{3},s_{2}}}}{\left( s_{3}+s_{2} \right)^{h_{15,s_{3},s_{2}}}+l_{15,s_{3},s_{2}}^{h_{15,s_{3},s_{2}}}}r_{15}$

$$\boldsymbol{x}_{\boldsymbol{j}}$$

| diagram | state | unit | description |
| --- | --- | --- | --- |
| 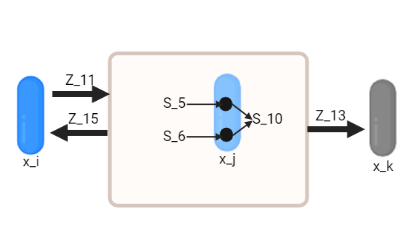 | $x_{j}$ | $\frac{{10}^{5}\text{cells}}{\mu L}$ | live Ri subpopulation with slow growth that consumes acetate ($s_{6}$) and/or lactate ($s_{5}$), and produces small amounts of butyrate ($s_{10}$) |

$$\frac{dx_{j}}{dt}=x_{j}\left[ \xi_{x_{j}}\mu max_{x_{j}}\left( \frac{s_{5}}{s_{5}+K_{x_{j},s_{5}}}+\frac{s_{6}}{s_{6}+K_{x_{i},s_{6}}} \right)-\left( Z_{15}+Z_{13} \right) \right]+x_{i}Z_{11}$$

$$\xi_{x_{j}}\left( pH \right)=\frac{1}{\varphi\left( p_{x_{j}};\alpha_{x_{j}},\frac{\alpha_{x_{j}}-1}{p_{x_{j}}} \right)}\varphi\left( pH;\alpha_{x_{j}},\frac{\alpha_{x_{j}}-1}{p_{x_{j}}} \right)$$

Fixed death rate:

$$Z_{\left( 13 \right)}=r_{13}$$

$$\boldsymbol{x}_{\boldsymbol{k}}$$

| diagram | state | unit | description |
| --- | --- | --- | --- |
| 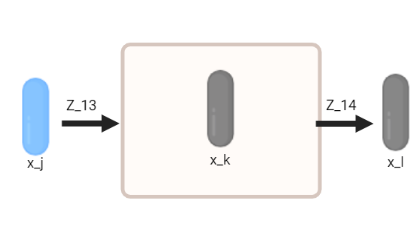 | $x_{k}$ | $\frac{{10}^{5}\text{cells}}{\mu L}$ | inactive Ri subpopulation that is (PI-positive) and eventually bursts |

$$\frac{dx_{k}}{dt}=x_{j}Z_{13}-x_{k}Z_{14}$$

$$\frac{dx_{l}}{dt}=x_{k}Z_{14}$$

Fixed burst rate:

$$Z_{14}=r_{14}$$

$$\boldsymbol{s}_{\boldsymbol{1}}$$

| diagram | state | unit | description |
| --- | --- | --- | --- |
| 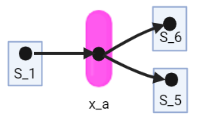 | $s_{1}$ | $mM$ | Trehalose concentration |

$$\frac{ds_{1}}{dt}=-\left( \gamma_{x_{a},s_{1}} \right)\left( \frac{s_{1}}{s_{1}+K_{x_{a},s_{1}}} \right)\left( \xi_{x_{a}}\mu max_{x_{a}}x_{a} \right)$$

$$\boldsymbol{s}_{\boldsymbol{2}}$$

| diagram | state | unit | description |
| --- | --- | --- | --- |
| 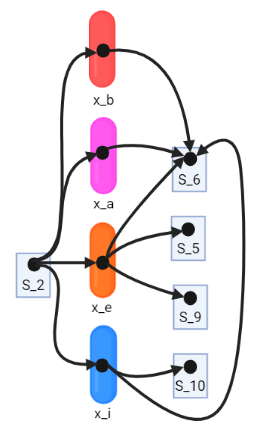 | $s_{2}$ | $mM$ | pyruvate concentration |

$\frac{ds_{2}}{dt}=-\left[ \left( \gamma_{x_{a},s_{2}} \right)\left( \frac{s_{2}}{s_{2}+K_{x_{a},s_{2}}} \right)\left( \xi_{x_{a}}\mu max_{x_{a}}x_{a} \right)+\left( \gamma_{x_{b},s_{2}} \right)\left( \frac{s_{2}}{s_{2}+K_{x_{b},s_{2}}} \right)\left( \xi_{x_{b}}\mu max_{x_{b}}x_{b} \right)+\left( \gamma_{x_{e},s_{2}} \right)\left( \frac{s_{2}}{s_{2}+K_{x_{e},s_{2}}} \right)\left( \xi_{x_{e}}\mu max_{x_{e}}x_{e} \right)+\left( \gamma_{x_{i},s_{2}} \right)\left( \frac{s_{2}}{s_{2}+K_{x_{i},s_{2}}} \right)\left( \xi_{x_{i}}\mu max_{x_{i}}x_{i} \right) \right]$

$$\boldsymbol{s}_{\boldsymbol{3}}$$

| Diagram | state | unit | description |
| --- | --- | --- | --- |
| 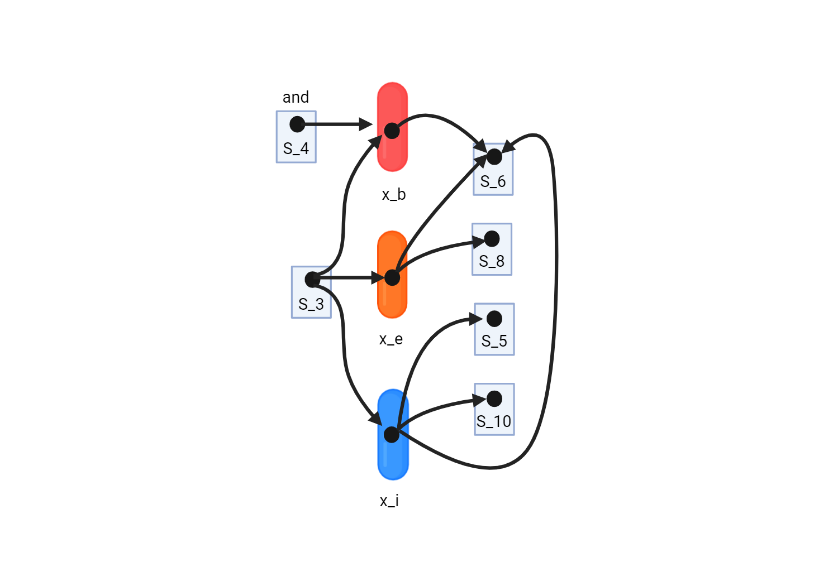 | $s_{3}$ | $mM$ | glucose concentration |

$$\frac{ds_{3}}{dt}=-\left[ \left( \gamma_{x_{b},s_{3}} \right)\left( \frac{s_{3}}{s_{3}+K_{x_{b},s_{3}}}\frac{s_{4}}{s_{4}+K_{x_{b},s_{4}}} \right)\left( \xi_{x_{b}}\mu max_{x_{b}}x_{b} \right)+\left( \gamma_{x_{e},s3} \right)\left( \frac{s_{3}}{s_{3}+K_{x_{e},s_{3}}} \right)\left( \xi_{x_{e}}\mu max_{x_{e}}x_{e} \right)+\left( \gamma_{x_{i},s_{3}} \right)\left( \frac{s_{3}}{s_{3}+K_{x_{i},s_{3}}} \right)\left( \xi_{x_{i}}\mu max_{x_{i}}x_{i} \right) \right]$$

$$\boldsymbol{s}_{\boldsymbol{4}}$$

| diagram | state | unit | description |
| --- | --- | --- | --- |
| 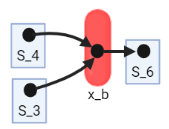 | $s_{4}$ | $mM$ | glutamate concentration |

$$\frac{ds_{4}}{dt}=-\left( \gamma_{x_{b},s_{4}} \right)\left( \frac{s_{3}}{s_{3}+K_{x_{b},s_{3}}}\frac{s_{4}}{s_{4}+K_{x_{b},s_{4}}} \right)\left( \xi_{x_{b}}\mu max_{x_{b}}x_{b} \right)$$

$$\boldsymbol{s}_{\boldsymbol{5}}$$

| Diagram | state | unit | description |
| --- | --- | --- | --- |
| 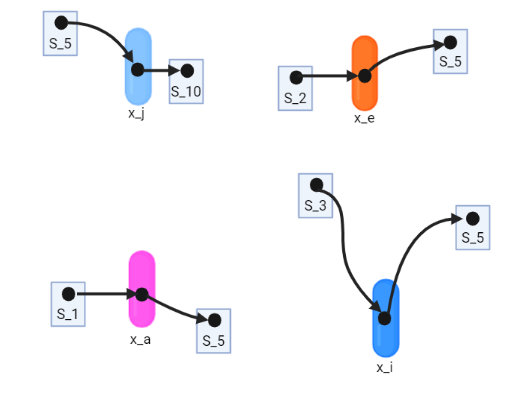 | $s_{5}$ | $mM$ | lactate concentration |

$$\frac{ds_{5}}{dt}=-\left( \gamma_{x_{j},s_{5}} \right)\left( \frac{s_{5}}{s_{5}+k_{x_{j},s_{5}}} \right)\left( \xi_{x_{j}}\mu max_{x_{j}}x_{j} \right)+\left( \gamma_{x_{a},s_{5},s_{1}} \right)\left( \frac{s_{1}}{s_{1}+k_{x_{a},s1}} \right)\left( \xi_{x_{a}}\mu max_{x_{a}}x_{a} \right)+\left( \gamma_{x_{e},s_{5},s_{2}} \right)\frac{s_{2}}{s_{2}+k_{x_{e},s_{2}}}\left( \xi_{x_{e}}\mu max_{x_{e}}x_{e} \right)+\left( \gamma_{x_{i},s_{5},s_{3}} \right)\frac{s_{3}}{s_{3}+K_{x_{i},s_{3}}}\left( \xi_{x_{i}}\mu max_{x_{i}}x_{i} \right)$$

$$\boldsymbol{s}_{\boldsymbol{6}}$$

| Diagram | state | unit | description |
| --- | --- | --- | --- |
| 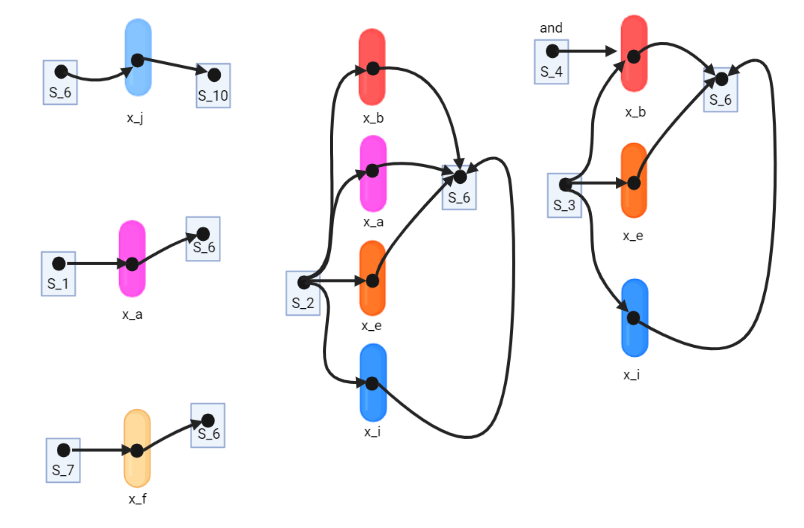 | $s_{6}$ | $mM$ | acetate concentration |

$$\frac{ds_{6}}{dt}=-\left( \gamma_{x_{j},s_{6}} \right)\left( \frac{s_{6}}{s_{6}+K_{x_{j},s_{6}}} \right)\left( \xi_{x_{j}}\mu max_{x_{j}}x_{j} \right)+\left[ \left( \gamma_{x_{a},s_{6},s_{1}} \right)\left( \frac{s_{1}}{s_{1}+K_{x_{a},s_{1}}} \right)+\left( \gamma_{x_{a},s_{6},s_{2}} \right)\left( \frac{s_{2}}{s_{2}+K_{x_{a},s_{2}}} \right) \right]\left( \xi_{x_{a}}\mu max_{x_{a}}x_{a} \right)+\left( \gamma_{x_{b},s_{6},s_{3},s_{4}} \right)\left( \frac{s_{3}}{s_{3}+K_{x_{b},s_{3}}}\frac{s_{4}}{s_{4}+K_{x_{b},s_{4}}} \right)\left( \xi_{x_{b}}\mu max_{x_{b}}x_{b} \right)+\left[ \left( \gamma_{x_{e},s_{6},s_{2}} \right)\left( \frac{s_{2}}{s_{2}+K_{x_{e},s_{2}}} \right)+\left( \gamma_{x_{e},s_{6},s_{3}} \right)\left( \frac{s_{3}}{s_{3}+K_{x_{e},s_{3}}} \right) \right]\left( \xi_{x_{e}}\mu max_{x_{e}}x_{e} \right)+\left( \gamma_{x_{f},s_{6},s_{7}} \right)\left( \frac{s_{7}}{s_{7}+K_{x_{f},s_{7}}} \right)\left( \xi_{x_{f}}\mu max_{x_{f}}x_{f} \right)+\left[ \left( \gamma_{x_{i},s_{6},s_{2}} \right)\left( \frac{s_{2}}{s_{2}+K_{x_{i},s_{2}}} \right)+\left( \gamma_{x_{i},s_{6},s_{3}} \right)\left( \frac{s_{3}}{s_{3}+K_{x_{i},s_{3}}} \right) \right]\left( \xi_{x_{i}}\mu max_{x_{i}}x_{i} \right)$$

$$\boldsymbol{s}_{\boldsymbol{7}}$$

| diagram | state | unit | description |
| --- | --- | --- | --- |
| 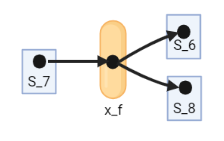 | $s_{7}$ | $mM$ | mannose concentration |

$$\frac{ds_{7}}{dt}=-\left( \gamma_{x_{f},s_{7}} \right)\left( \frac{s_{7}}{s_{7}+K_{x_{f},s7}} \right)\left( \xi_{x_{f}}\mu max_{x_{f}}x_{f} \right)$$

$$\boldsymbol{s}_{\boldsymbol{8}}$$

| diagram | state | unit | description |
| --- | --- | --- | --- |
| 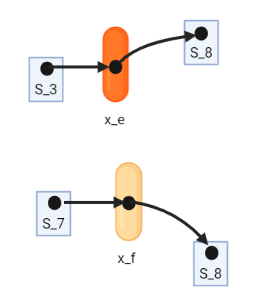 | $s_{8}$ | $mM$ | succinate concentration |

$\frac{ds_{8}}{dt}=\left( \gamma_{x_{e},s_{8},s_{3}} \right)\left( \frac{s_{3}}{s_{3}+K_{x_{e},s_{3}}} \right)\left( \xi_{x_{e}}\mu max_{x_{e}}x_{e} \right)+\left( \gamma_{x_{f},s_{8},s_{7}} \right)\left( \frac{s_{7}}{s_{7}+K_{x_{f},s_{7}}} \right)\left( \xi_{x_{f}}\mu max_{x_{f}}x_{f} \right)$

$$\boldsymbol{s}_{\boldsymbol{9}}$$

| Diagram | state | unit | description |
| --- | --- | --- | --- |
| 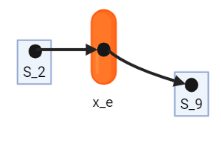 | $s_{9}$ | $mM$ | formate concentration |

$$\frac{ds_{9}}{dt}=\left( \gamma_{x_{e},s_{9},s_{2}} \right)\left( \frac{s_{2}}{s_{2}+K_{x_{e},s_{2}}} \right)\left( \xi_{x_{e}}\mu max_{x_{e}}x_{e} \right)$$

$$\boldsymbol{s}_{\boldsymbol{10}}$$

| diagram | state | unit | description |
| --- | --- | --- | --- |
| 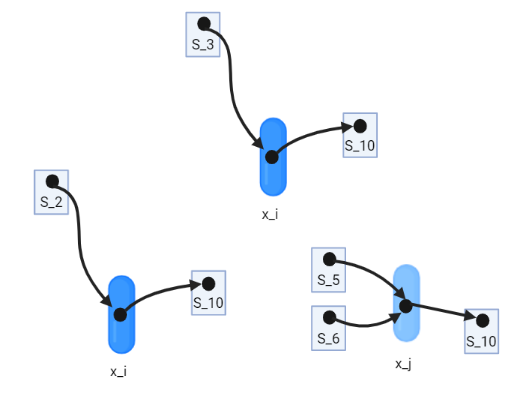 | $s_{10}$ | $mM$ | butyrate concentration |

$\frac{ds_{10}}{dt}=\left[ \left( \gamma_{x_{i},s_{10},s_{2}} \right)\left( \frac{s_{2}}{s_{2}+K_{x_{i},s_{2}}} \right)+\left( \gamma_{x_{i},s_{10},s_{3}} \right)\left( \frac{s_{3}}{s_{3}+K_{x_{i},s_{3}}} \right) \right]\left( \xi_{x_{i}}\mu max_{x_{i}}x_{i} \right)+\left[ \left( \gamma_{x_{j},s_{10},s_{5}} \right)\left( \frac{s_{5}}{s_{5}+K_{x_{j},s_{5}}} \right)+\left( \gamma_{x_{j},s_{10},s_{6}} \right)\left( \frac{s_{6}}{s_{6}+K_{x_{j},s_{6}}} \right) \right]\left( \xi_{x_{j}}\mu max_{x_{j}}x_{j} \right)$

| Parameters  # | species | Parameter | symbol | fitted values | description |
| --- | --- | --- | --- | --- | --- |
| 1 | Bh | xa_mumax | $\mu max_{x_{a}}$ | $0.192$ | maximum growth rate of $x_{a}$ |
| 2 | Bh | xb_mumax | $\mu max_{x_{b}}$ | $0.978$ | maximum growth rate of $x_{b}$ |
| 3 | Bh | xa_pHopt | $p_{x_{a}}$ | $6.818$ | $x_{a}$ optimal pH |
| 4 | Bh | xb_pHopt | $p_{x_{b}}$ | $6.539$ | $x_{b}$ optimal pH |
| 5 | Bh | xa_pHalpha | $\alpha_{x_{a}}$ | $46.273$ | $x_{a}$ pH sensitivity |
| 6 | Bh | xb_pHalpha | $\alpha_{x_{b}}$ | $62.518$ | $x_{b}$ pH sensitivity |
| 7 | Bh | xa_k_s1 | $K_{x_{a},s_{1}}$ | $0.287$ | Monod constant for trehalose ($s_{1}$) consumption by $x_{a}$ |
| 8 | Bh | xa_k_s2 | $K_{x_{a},s_{2}}$ | $2.391$ | Monod constant for pyruvate ($s_{2}$) consumption by $x_{a}$ |
| 9 | Bh | xb_k_s3 | $K_{x_{b},s_{3}}$ | $0.103$ | Monod constant for glucose ($s_{3}$) consumption by $x_{b}$ |
| 10 | Bh | xb_k_s4 | $K_{x_{b},s_{4}}$ | $1.270$ | Monod constant for glutamate ($s_{4}$) consumption by $x_{b}$ |
| 11 | Bh | xb_k_s2 | $K_{x_{b},s1}$ | $0.500$ | Monod constant for pyruvate ($s_{2}$) consumption by $x_{b}$ |
| 12 | Bh | xa_g_s1 | $\gamma_{x_{a},s_{1}}$ | $1.241$ | stoichiometric constant for trehalose ($s_{1}$) consumption by $x_{a}$ |
| 13 | Bh | xa_g_s2 | $\gamma_{x_{a},s_{2}}$ | $1.000$ | stoichiometric constant for pyruvate ($s_{2}$) consumption by $x_{a}$ |
| 14 | Bh | xa_g_s6_s1 | $\gamma_{x_{a},s_{6},s_{1}}$ | $-1.596$ | stoichiometric constant for acetate ($s_{6}$) production from $x_{a}$ consuming trehalose ($s_{1}$) |
| 15 | Bh | xa_g_s5_s1 | $\gamma_{x_{a},s_{5},s_{1}}$ | $-2.999$ | stoichiometric constant for lactate ($s_{5}$) production from $x_{a}$ consuming trehalose ($s_{1}$) |
| 16 | Bh | xa_g_s6_s2 | $\gamma_{x_{a},s_{6},s_{2}}$ | $-1.869$ | stoichiometric constant for acetate ($s_{6}$) production from $x_{a}$ consuming pyruvate ($s_{2}$) |
| 17 | Bh | xb_g_s3 | $\gamma_{x_{b},s_{3}}$ | $3.763$ | stoichiometric constant for glucose ($s_{3}$) consumption by $x_{b}$ |
| 18 | Bh | xb_g_s4 | $\gamma_{x_{b},s_{4}}$ | $0.500$ | stoichiometric constant for glutamate ($s_{4}$) consumption by $x_{b}$ |
| 19 | Bh | xb_g_s2 | $\gamma_{x_{b},s_{2}}$ | $10.000$ | stoichiometric constant for pyruvate ($s_{2}$) consumption by $x_{b}$ |
| 20 | Bh | xb_g_s6_s3_s4 | $\gamma_{x_{b},s_{6},s_{3},s_{4}}$ | $-2.999$ | stoichiometric constant for acetate ($s_{6}$) production from $x_{b}$ consuming glucose ($s_{3}$) and glutamate ($s_{4}$) |
| 21 | Bh | xb_g_s6_s2 | $\gamma_{x_{b},s_{6},s_{2}}$ | $-4.999$ | stoichiometric constant for acetate ($s_{6}$) production from $x_{b}$ consuming pyruvate ($s_{2}$) |
| 22 | Bh | z1_r | $r_{1}$ | $0.025$ | rate of transition from $x_{a}$ to $x_{b}$ |
| 23 | Bh | z2_r | $r_{2}$ | $1.5$ | rate of transition from $x_{b}$ to $x_{a}$ |
| 24 | Bh | z3_r | $r_{3}$ | $0.00001$ | rate of transition from $x_{a}$ to $x_{c}$ |
| 25 | Bh | z4_r | $r_{4}$ | $0.296$ | rate of transition from $x_{b}$ to $x_{c}$ |
| 26 | Bh | z5_r | $r_{5}$ | $0.035$ | rate of transition from $x_{c}$ to $x_{d}$ |
| 27 | Bh | z1_l_s1 | $l_{1,s_{1}}$ | $1e-07$ | half saturation constant for trehalose ($s_{1}$) |
| 28 | Bh | z2_l_s1 | $l_{2,s1}$ | $0.1000$ | half saturation constant for trehalose ($s_{1}$) |
| 29 | Bh | z4_l_s3_s4 | $l_{3,s_{3},s_{4}}$ | $0.0001$ | half saturation constant for glucose ($s_{3}$) and glutamate ($s_{4}$) |
| 30 | Bh | z1_h_s1 | $h_{1,s_{1}}$ | $0.999$ | Hill coefficient for trehalose ($s_{1}$) |
| 31 | Bh | z2_h_s1 | $h_{2,s_{1}}$ | $10.0000$ | Hill coefficient for trehalose ($s_{1}$) |
| 32 | Bh | z4_h_s3_s4 | $h_{3,s_{3},s_{4}}$ | $2.905$ | Hill coefficient for glucose ($s_{3}$) and glutamate ($s_{4}$) |
| 33 | Bt | xe_mumax | $\mu max_{x_{e}}$ | $0.921$ | maximum growth rate of $x_{e}$ |
| 34 | Bt | xf_mumax | $\mu max_{x_{f}}$ | $1.196$ | maximum growth rate of $x_{f}$ |
| 35 | Bt | xe_pHopt | $p_{x_{e}}$ | $7.507$ | $x_{e}$ optimal pH |
| 36 | Bt | xf_pHopt | $p_{x_{f}}$ | $6.984$ | $x_{f}$ optimal pH |
| 37 | Bt | xe_pHalpha | $\alpha_{x_{e}}$ | $59.823$ | $x_{e}$ pH sensitivity |
| 38 | Bt | xf_pHalpha | $\alpha_{x_{f}}$ | $71.816$ | $x_{f}$ pH sensitivity |
| 39 | Bt | xe_k_s3 | $K_{x_{e},s_{3}}$ | $0.353$ | Monod constant for glucose ($s_{3}$) consumption by $x_{e}$ |
| 40 | Bt | xe_k_s2 | $K_{x_{e},s_{2}}$ | $9.999$ | Monod constant for pyruvate ($s_{2}$) consumption by $x_{e}$ |
| 41 | Bt | xf_k_s7 | $K_{x_{f},s_{7}}$ | $0.105$ | Monod constant for mannose ($s_{7}$) consumption by $x_{f}$ |
| 42 | Bt | xe_g_s2 | $\gamma_{x_{e},s_{2}}$ | $2.901$ | stoichiometric constant for pyruvate ($s_{2}$) consumption by $x_{e}$ |
| 43 | Bt | xe_g_s3 | $\gamma_{x_{e},s_{3}}$ | $1.026$ | stoichiometric constant for glucose ($s_{3}$) consumption by $x_{e}$ |
| 44 | Bt | xe_g_s5_s2 | $\gamma_{x_{e},s_{5},s_{2}}$ | $-1.718$ | stoichiometric constant for lactate ($s_{5}$) production from $x_{e}$ consuming pyruvate ($s_{2}$) |
| 45 | Bt | xe_g_s6_s2 | $\gamma_{x_{e},s_{6},s_{2}}$ | $-0.260$ | stoichiometric constant for acetate ($s_{6}$) production from $x_{e}$ consuming pyruvate ($s_{2}$) |
| 46 | Bt | xe_g_s6_s3 | $\gamma_{x_{e},s_{6},s_{3}}$ | $-0.996$ | stoichiometric constant for acetate ($s_{6}$) production from $x_{e}$ consuming glucose ($s_{3}$) |
| 47 | Bt | xe_g_s8_s3 | $\gamma_{x_{e},s_{8},s_{3}}$ | $-0.625$ | stoichiometric constant for succinate ($s_{8}$) production by $x_{e}$ consuming glucose ($s_{3}$) |
| 48 | Bt | xe_g_s9_s2 | $\gamma_{x_{e},s_{9},s_{2}}$ | $-0.615$ | stoichiometric constant for formte ($s_{9}$) production by $x_{e}$ consuming pyruvate ($s_{2}$) |
| 49 | Bt | xf_g_s7 | $\gamma_{x_{f},s_{7}}$ | $0.404$ | stoichiometric constant for mannose ($s_{7}$) consumption by $x_{f}$ |
| 50 | Bt | xf_g_s6_s7 | $\gamma_{x_{f},s_{6},s_{7}}$ | $-0.853$ | stoichiometric constant for acetate ($s_{6}$) prouduction from $x_{f}$ consuming mannose ($s_{7}$) |
| 51 | Bt | xf_g_s8_s7 | $\gamma_{x_{f},s_{8},s_{7}}$ | $-2.025$ | stoichiometric constant for succinate ($s_{8}$) production by $x_{f}$ consuming mannose ($s_{7}$) |
| 52 | Bt | z6_r | $r_{6}$ | $1.495$ | rate of transition from $x_{e}$ to $x_{f}$ |
| 53 | Bt | z7_r | $r_{7}$ | $0.957$ | rate of transition from $x_{e}$ to $x_{g}$ |
| 54 | Bt | z8_r | $r_{8}$ | $0.076$ | rate of transition from $x_{f}$ to $x_{g}$ |
| 55 | Bt | z9_r | $r_{9}$ | $0.004$ | rate of transition from $x_{g}$ to $x_{h}$ |
| 56 | Bt | z10_r | $r_{10}$ | $0.0001$ | rate of transition from $x_{f}$ to $x_{e}$ |
| 57 | Bt | z6_l_s3 | $l_{6,s_{3}}$ | $0.009$ | half-saturation constant for glucose ($s_{3}$) |
| 58 | Bt | z6_l_s7 | $l_{6,s_{7}}$ | $0.509$ | half-saturation constant for mannose ($s_{7}$) |
| 59 | Bt | z7_l_s3 | $l_{7,s_{3}}$ | $0.006$ | half-saturation constant for glucose ($s_{3}$) |
| 60 | Bt | z7_l_pH | $l_{7,\text{pH}}$ | $7.162$ | half-saturation constant for the pH |
| 61 | Bt | z8_l_s7 | $l_{8,s7}$ | $1.000$ | half-saturation constant for mannose ($s_{7}$) |
| 62 | Bt | z8_l_pH | $l_{8,\text{pH}}$ | $6.000$ | half-saturation constant for the pH |
| 63 | Bt | z10_l_s3 | $l_{10,s_{3}}$ | $0.500$ | half-saturation constant for glucose ($s_{3}$) |
| 64 | Bt | z6_h_s3 | $h_{6,s_{3}}$ | $1.002$ | Hill coefficient for glucose ($s_{3}$) |
| 65 | Bt | z6_h_s7 | $h_{6,s_{7}}$ | $1.002$ | Hill coeffiecient for mannose ($s_{7}$) |
| 66 | Bt | z7_h_s3 | $h_{7,s_{3}}$ | $1.459$ | Hill coefficient for glucose ($s_{3}$) |
| 67 | Bt | z7_h_pH | $h_{7,\text{pH}}$ | $10.00$ | Hill coefficient for the pH |
| 68 | Bt | z8_h_s7 | $h_{8,s7}$ | $29.999$ | Hill coefficient for mannose ($s_{7}$) |
| 69 | Bt | z8_h_pH | $h_{8,pH}$ | $10.000$ | Hill coefficient for the pH |
| 70 | Bt | z10_h_s7 | $h_{10,s_{7}}$ | $10.0000$ | Hill coefficient for mannose ($s_{7}$) |
| 71 | Ri | xi_mumax | $\mu max_{x_{i}}$ | $0.705$ | maximum growth rate of $x_{i}$ |
| 72 | Ri | xj_mumax | $\mu max_{x_{j}}$ | $0.015$ | maximum growth rate of $x_{j}$ |
| 73 | Ri | xi_pHopt | $p_{xi}$ | $7.768$ | $x_{i}$ optimal pH |
| 74 | Ri | xj_pHopt | $p_{x_{j}}$ | $7.624$ | $x_{j}$ optimal pH |
| 75 | Ri | xi_pHalpha | $\alpha_{x_{i}}$ | $39.525$ | $x_{i}$ pH sensitivity |
| 76 | Ri | xj_pHalpha | $\alpha_{x_{j}}$ | $19.999$ | $x_{j}$ pH sensitivity |
| 77 | Ri | xi_k_s2 | $K_{x_{i},s_{2}}$ | $0.547$ | Monod constant for pyruvate ($s_{2}$) by $x_{i}$ |
| 78 | Ri | xi_k_s3 | $K_{x_{i},s_{3}}$ | $6.865$ | Monod constant for glucose ($s_{3}$) by $x_{i}$ |
| 79 | Ri | xj_k_s5 | $K_{x_{j},s_{5}}$ | $9.999$ | Monod constant for lactate ($s_{5}$) by $x_{j}$ |
| 80 | Ri | xj_k_s6 | $K_{x_{j},s_{6}}$ | $0.275$ | Monod constant for acetate ($s_{6}$) by $x_{j}$ |
| 81 | Ri | xi_g_s2 | $\gamma_{xi,s2}$ | $2.783$ | stoichiometric constant for pyruvate ($s_{2}$) consumption by $x_{i}$ |
| 82 | Ri | xi_g_s3 | $\gamma_{x_{i},s_{3}}$ | $1.759$ | stoichiometric constant for glucose ($s_{3}$) consumption by $x_{i}$ |
| 83 | Ri | xj_g_s5 | $\gamma_{x_{j},s_{5}}$ | $1.960$ | stoichiometric constant for lactate ($s_{5}$) consumption by $x_{j}$ |
| 84 | Ri | xj_g_s6 | $\gamma_{x_{j},s_{6}}$ | $0.853$ | stoichiometric constant for acetate ($s_{6}$) consumption by $x_{j}$ |
| 85 | Ri | xi_g_s5_s3 | $\gamma_{x_{i},s_{5},s_{e}}$ | $-0.377$ | stoichiometric constant for lactate ($s_{5}$) production from $x_{i}$ consuming glucose ($s_{3}$) |
| 86 | Ri | xi_g_s6_s2 | $\gamma_{x_{i},s_{6},s_{2}}$ | $-0.305$ | stoichiometric constant for acetate ($s_{6}$) production from $x_{i}$ consuming pyruvate ($s_{2}$) |
| 87 | Ri | xi_g_s6_s3 | $\gamma_{x_{i},s_{6},s_{3}}$ | $-0.727$ | stoichiometric constant for acetate ($s_{6}$) production from $x_{i}$ consuming glucose ($s_{3}$) |
| 88 | Ri | xi_g_s10_s2 | $\gamma_{x_{i},s_{10},s_{2}}$ | $-1.994$ | stoichiometric constant for butyrate ($s_{10}$) production by $x_{i}$ consuming pyruvate ($s_{2}$) |
| 89 | Ri | xi_g_s10_s3 | $\gamma_{x_{i},s_{10},s_{3}}$ | $-6.865$ | stoichiometric constant for butyrate ($s_{10}$) production by $x_{i}$ consuming glucose ($s_{3}$) |
| 90 | Ri | xj_g_s10_s5 | $\gamma_{x_{j},s_{10},s_{5}}$ | $-2.988$ | stoichiometric constant for butyrate ($s_{10}$) production by $x_{j}$ consuming lactate ($s_{5}$) |
| 91 | Ri | xj_g_s10_s6 | $\gamma_{x_{j},s_{10},s_{6}}$ | $-1.994$ | stoichiometric constant for butyrate ($s_{10}$) production by $x_{j}$ consuming acetate ($s_{6}$) |
| 92 | Ri | z11_r | $r_{11}$ | $0.021$ | rate of transition from $x_{i}$ to $x_{j}$ |
| 93 | Ri | z12_r | $r_{12}$ | $1.999$ | rate of transition from $x_{i}$ to $x_{l}$ |
| 94 | Ri | z13_r | $r_{13}$ | $0.009$ | rate of transition from $x_{j}$ to $x_{k}$ |
| 95 | Ri | z14_r | $r_{14}$ | $0.00001$ | rate of transition from $x_{k}$ to $x_{l}$ |
| 96 | Ri | Z15_r | $r_{15}$ | $0.214$ |  |
| 97 | Ri | z11_l_s5 | $l_{11,s_{5}}$ | $2.331$ | half-saturation constant for lactate ($s_{5}$) |
| 98 | Ri | z12_l_s3_s2 | $l_{12,s_{3},s_{2}}$ | $0.003$ | half-saturation constant for glucose ($s_{3}$) and pyruvate ($s_{2}$) |
| 99 | Ri | z15_l_s3_s2 | $l_{15,s_{3},s_{2}}$ | $0.500$ | half-saturation constant for glucose ($s_{3}$) and pyruvate ($s_{2}$) |
| 100 | Ri | z11_h_s5 | $h_{9,s_{5}}$ | $29.999$ | Hill coefficient for lactate ($s_{5}$) |
| 101 | Ri | z12_h_s3_s2 | $h_{10,s3_{s}2}$ | $1.112$ | Hill coefficient for glucose ($s_{3}$) and pyruvate ($s_{2}$) |
| 102 | Ri | z15_h_s3_s2 | $h_{15,s_{3},s_{2}}$ | $10.000$ | Hill coefficient for glucose ($s_{3}$) and pyruvate ($s_{2}$) |

Finding model parameters

We used batch monoculture experiments to estimate the parameters for the state equations in our model. The codes that execute the complete procedure described below for each of the strains are available [here](https://github.com/danielriosgarza/hungerGamesModel/tree/main/scripts/parameterFIt).

*Moving average*

We carried out three sets of monoculture measurements for cell counts, metabolite concentrations, and pH, with each set including at least two replicates. These experiments, conducted under nearly identical conditions, showed consistent kinetics across different days, despite some variability (e.g. lactate). We averaged measurements within regular intervals to establish a consensus kinetic curve for each state, which was used to estimate the model's kinetic parameters.

For example, the black traced line below shows the moving average:

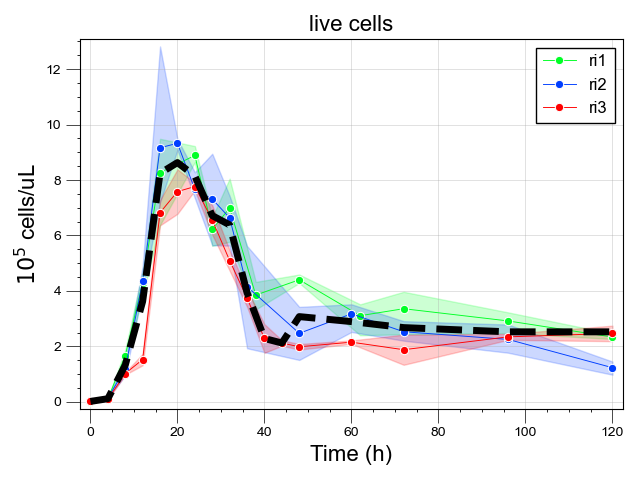

*Cubic splines*

Moving averages were interpolated by cubic-splines, a method for smoothing data. Cubic spline interpolation is a mathematical technique often employed in data analysis to create a smooth curve that intersects a given set of points (in this case, the moving averages). This technique involves constructing a series of piecewise continuous polynomials of degree three that fit the data.

Cubic splines provide the advantage of calculating state values at any given point in time, not just at the sampled intervals. This means we can evaluate the state at any arbitrary point within the range of the measured time-points, not merely at specified time intervals. Using cubic splines, we can directly compare the smoothed values from the data (the spline's function values) with the solutions of our model's differential equations. This comparison is accomplished by using the simulation times from the model as inputs for the cubic spline functions.

For example, the orange line is the spline:

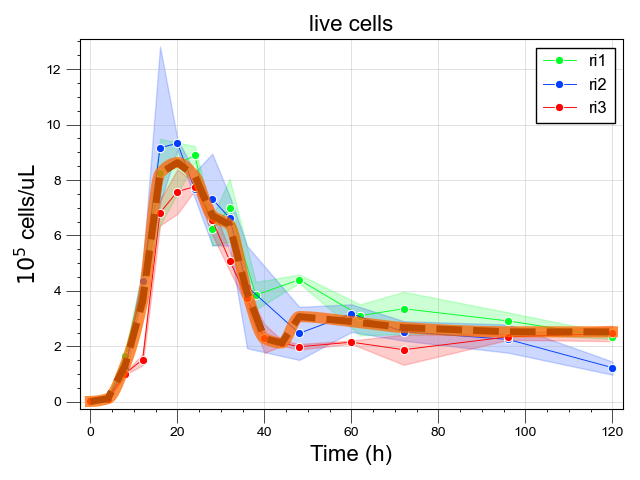

*Optimization*

Finally, we searched for the parameter set that minimizes the difference between the [states](https://github.com/danielriosgarza/hungerGamesModel/wiki/State-equations#states) in our model simulations and their corresponding smoothed moving averages. To compute the discrepancy between the model simulations and the experimental data, we used the Hubber loss function, which is a function that is robust against eventual outliers. We then minimized this loss function using Powell's method, an approach that approximates the function using a quadratic model. The method sequentially minimizes this approximation along one direction at a time, generating a set of mutually conjugate search directions to navigate towards the minimum. The parameters that minimize this loss were subsequently used in all downstream simulations of our model.

In summary, the process of fitting our model to sets of experimental data involved the following steps:

Acquiring the moving average from experimental data that represent our model states

Fitting cubic splines to these averages to enable a smooth estimation of their values within their measured time frames;

Adjusting model parameters by minimizing the Huber loss function between model simulations and the smoothed moving averages (simulated by the cubic splines), using Powell's optimization method.
