## Supplementary Text S2 for "Discovery of alternative stable states in a synthetic human gut microbial community"

**Phenomenological model**

Simulations of the detailed mechanistic model described in Supplementary Text S1 suggested a conceptual mechanism for the emergence of alternative community states (See Fig. 3 of the main text):

- Species genomes encode alternative phenotypes.
- Subpopulations executing specific metabolic programs emerge triggered by environment cues.
- Subpopulations may occupy different niches, compete for different resources, and have different physiological capacities, even if belonging to the same species.
- A key factor is the ecological interactions between subpopulations. When a subpopulation is a strong competitor with other subpopulations, the community state can be significantly different when this subpopulation is highly expressed, compared to when it is inhibited.

We implemented this general mechanism in a simple generalized Lotka-Volterra model, first for our three species with parameters qualitatively reflecting the parameters of the mechanistic models, then we built a simulation of a large community, which shows that a super competitor phenotype (such as *Blautia hydrogenotrophica*’s glucose consuming phenotype in our system) could drive the system towards alternative states. Here, we give details on the implementation.

Below we provide an overview of the equations used in this phenomenological model. For a detailed computational implementation, refer to the online repository: <https://github.com/danielriosgarza/hungerGamesModel>

**Generic model**

Species are characterized by one or more subpopulations, each with a growth rate, an interaction vector, and incoming and outgoing flow to subpopulations of the same species, but with different phenotypes. The transitions followed a similar implementation as the ones in the mechanistic model (refer to Supplementary Text S1, “Subpopulations” section), consisting of a response ($Z$) of three types to a single environment cue ($e)$: independent, activation, or inhibition.

Independent:

$$Z=1$$

Activation:

$$Z=\frac{e^{h}}{K^{h}+e^{h}}$$

Inhibition:

$$Z=\frac{K^{h}}{K^{h}+e^{h}}$$

The general growth equation of the i^th^ subpopulation is given by:

$$\frac{dN_{i}}{dt}=N_{i}\left( \mu_{i}+\sum_{k} \alpha_{ij}N_{k} \right)+\sum Z_{in}r_{in}N_{in}-\sum Z_{out}r_{out}N_{out}$$

Where $i$ and $k$ are subpopulation indices, $N$s are subpopulation concentrations, $\mu$ is the growth rate, $\alpha s$ are interactions, and $Z_{in}r_{in}N_{in}$ are sources of incoming subpopulations from the same species and $Z_{out}r_{out}N_{out}$ are sinks of outgoing subpopulation from the same species.

**Three-species model**

The three species system shown in main Fig.4 was defined by the following six differential equations (the environment cue is referred to as $e$):

*Blautia hydrogenotrophica* trehalose phenotype ($x_{a}$):

$$\frac{dx_{a}}{dt}=x_{a}(\mu_{x_{a}}-x_{a}-f_{1}r_{1})+x_{b}f_{2}r_{2}$$

$$f_{1}=\frac{e^{h_{1}}}{K_{1}^{h_{1}}+e^{h_{1}}}$$

$$f_{2}=\frac{K_{2}^{h_{2}}}{K_{2}^{h_{2}}+e^{h_{2}}}$$

*Blautia hydrogenotrophica* glucose phenotype ($x_{b}$):

$$\frac{dx_{b}}{dt}=x_{b}\left( \mu_{x_{b}}-x_{b}-0.4x_{e}-0.3x_{i}-f_{2}r_{2} \right)+x_{a}f_{1}r_{1}$$

*Bacteroides thetaiotaomicron* glucose phenotype ($x_{e}$) :

$$\frac{dx_{e}}{dt}=x_{e}\left( \mu_{x_{e}}-0.9x_{b}-x_{e}-0.3x_{i}-f_{3}r_{3} \right)+x_{f}f_{4}r_{4}$$

$$f_{3}=f_{4}=1$$

Transitions are independent of $\epsilon$

*Bacteroides thetaiotaomicron* mannose phenotype ($x_{f}$):

$$\frac{dx_{f}}{dt}=x_{f}\left( \mu_{x_{f}}-0.4x_{b}-x_{f}-f_{4}r_{4} \right)+x_{e}f_{3}r_{3}$$

*Roseburia intestinalis* fast growth mode ($x_{i}$):

$$\frac{dx_{i}}{dt}=x_{i}\left( \mu_{x_{i}}-0.9x_{b}-0.4x_{e}-x_{i}-f_{5}r_{5} \right)+x_{j}f_{6}r_{6}$$

$$f_{5}=\frac{e^{h_{5}}}{K_{5}^{h_{5}}+e^{h_{5}}}$$

$$f_{6}=\frac{e^{h_{6}}}{K_{6}^{h_{6}}+e^{h_{6}}}$$

*Roseburia intestinalis* slow growth mode:

$$\frac{dx_{j}}{dt}=x_{j}\left( \mu_{x_{j}}+0.1\left[ x_{a}+x_{b}+x_{e}+x_{f}+x_{i} \right]-x_{j}-f_{6}r_{6} \right)+x_{i}f_{5}r_{5}$$

**Parameters**

| **Parameter** | **value** |
| --- | --- |
| $\mu_{x_{a}}$ | 0.192 |
| $\mu_{x_{b}}$ | 0.978 |
| $\mu_{x_{e}}$ | 0.921 |
| $\mu_{x_{f}}$ | 1.190 |
| $\mu_{x_{i}}$ | 0.705 |
| $\mu_{x_{j}}$ | 0.010 |
| $r_{1}$ | 0.025 |
| $r_{2}$ | 0.850 |
| $r_{3}$ | 0.495 |
| $r_{4}$ | 0.001 |
| $r_{5}$ | 0.021 |
| $r_{6}$ | 0.214 |
| $h_{1}$ | 9.000 |
| $h_{2}$ | 10.000 |
| $h_{5}$ | 9.000 |
| $h_{6}$ | 10.000 |
| $K_{1}$ | 0.100 |
| $K_{2}$ | 0.100 |
| $K_{5}$ | 0.100 |
| $K_{6}$ | 0.100 |

**Large random system**

To simulate fifty species with one bacterium that switches phenotype in response to an environmental cue, we generated random growth rates from a uniform distribution and random interaction vectors from a beta distribution, transformed to assume values between -1 and 1. Next, we skewed the distribution towards negative interactions by sampling from larger alpha parameters (the resulting distributions are shown in Fig. 4C). We then performed 1,000 simulations using the stochastic Stratonovish Heun integrator from the sdeint package (https://github.com/mattja/sdeint) with a fixed diffusion parameter, which simulates the addition of white noise during each simulation (Brownian motion). The environment cue parameter (e) was sampled from a uniform distribution. The simulations shown in Fig.4 were performed with the following parameters:

- Beta distribution from background community: $\alpha= \beta= 51$
- $\alpha$ parameters for the strongly interacting phenotype, respectively: 51, 68, and 85
- Diffusion parameter for the stochastic ODE: 0.2
- Growth rate of weakly interacting phenotype: 0.3
- Growth rate of strongly interacting phenotype: 0.5
- Activation rate of the transition function: 0.1
- Inhibition rate of the transition function: 0.5
- Hill coefficients: 10
- Halfmax constants: 0.1
