## Supplementary figures and images for "Discovery of alternative stable states in a synthetic human gut microbial community"

### Supplementary Movie S1

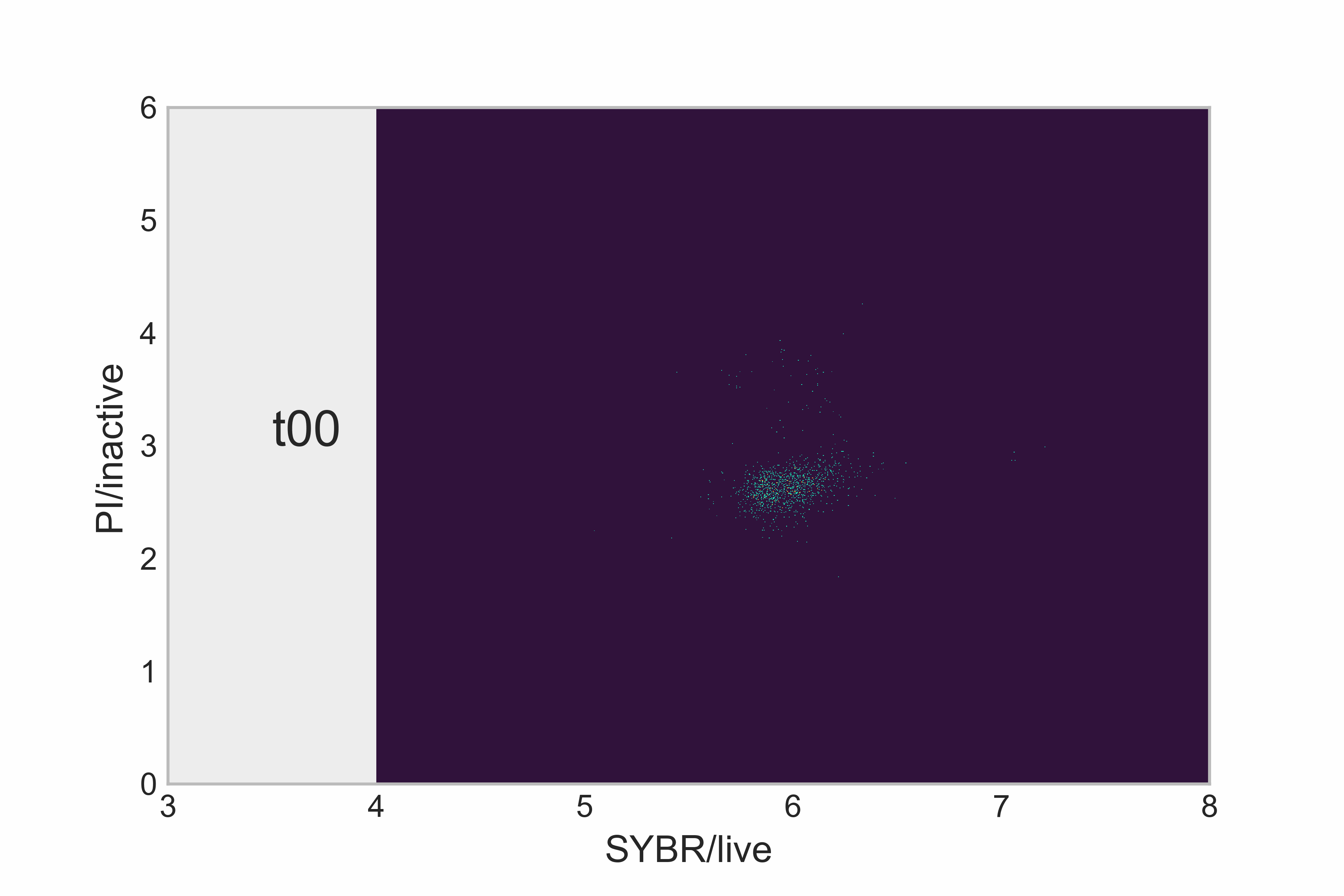
